# Autoregulation of switching behavior by cellular compartment size

**DOI:** 10.1101/2021.10.01.462711

**Authors:** Monika Jozsa, Tihol Ivanov Donchev, Rodolphe Sepulchre, Timothy O’Leary

## Abstract

Many kinds of cellular compartments comprise decision making mechanisms that control growth and shrinkage of the compartment in response to external signals. Key examples include synaptic plasticity mechanisms that regulate the size and strength of synapses in the nervous system. However, when synaptic compartments and postsynaptic densities are small such mechanisms operate in a regime where chemical reactions are discrete and stochastic due to low copy numbers of the species involved. In this regime, fluctuations are large relative to mean concentrations, and inherent discreteness leads to breakdown of mass action kinetics. Understanding how synapses and other small compartments achieve reliable switching in the low copy number limit thus remains a key open problem. We propose a novel self regulating signaling motif that exploits the breakdown of mass action kinetics to generate a reliable size-regulated switch. We demonstrate this in simple two and three-species chemical reaction systems and uncover a key role for inhibitory loops among species in generating switching behavior. This provides an elementary motif that could allow size dependent regulation in more complex reaction pathways and may explain discrepant experimental results on well-studied biochemical pathways.

## Introduction

A switch is a fundamental operation required of biochemical signaling. In essence, a switch is a system that reliably transitions between at least two distinct states while exhibiting a memory, or hysteresis, that preserves the state in the absence of input. Such systems are critical for cellular decision-making processes, including apoptosis (1–3), cell fate decisions (4–6) and synaptic plasticity (7). Much of what we know about switching behavior in these contexts relies on mass-action kinetics that model concentrations of biochemical species as continuous, deterministic quantities (1, 2, 5, 8–11).

However, many cellular compartments, including synaptic spines and postsynaptic densities, contain concentrations of key signaling enzymes that correspond to hundreds or even tens of individual molecules (12, 13), where mass-action kinetics break down. Moreover, even relatively simple biochemical reaction motifs exhibit qualitative changes in behavior as system size transitions between the microscopic and macroscopic limit. A famous example includes the so-called non-cooperative toggle switch motif (14), a pair of mutually inhibitory enzymes which, under broad parameter regimes, acquires a single, stable equilibrium in the mass-action limit that transitions into two distinct stochastic modes as system size decreases (15, 16). This system thus acts as a switch in the small system size limit but has a single stable equilibrium as the system grows.

In this paper we put forward the hypothesis that biological systems may exploit the qualitative transition between discrete, stochastic dynamics at the microscopic scale, and deterministic, continuous dynamics at the macroscopic limit. Many cellular compartments grow in size, and the growth process itself depends on the outcome of a cellular decision: a switching event. This raises the possibility that system size can act as a feedback signal to self regulate switching behavior.

Using analysis and simulations of simple reaction motifs, we substantiate this hypothesis with a novel working model. We show that generic, mutually inhibitory reactions between two or more species can give rise to switch-like behavior in the microscopic limit that gives way to a stable behavior as the system size increases. Furthermore, these systems can regulate the size evolution of small systems using their switch-like behavior, maintain the stability of large systems, and integrate external input that encourages growth.

Our results and hypothesis provide a new signaling motif that may be exploited by cellular systems to make growth decisions in a regime where the inherent randomness and discreteness of biochemical signals governs the dynamics.

### A size-dependent switch motif

We begin by exploring simple reaction networks that exhibit size-dependent bistability: the system behaves as a switch when the total number of molecules is low (in the tens) and has a single stable equilibrium as system size increases. Our goal is not to model any specific reaction pathway in detail, but to characterize minimal motifs of two or three reacting species that can support switching behavior. Such motifs may constitute building blocks or sub-networks that confer switching functionality to more complex chemical reaction networks, including those governing synaptic plasticity and growth of synaptic densities (12).

A well-studied motif (14–17), shown in Figure 1 (a), is a pair of mutually inhibitory species, *x*_1_ and *x*_2_, described by the following reaction scheme (where *x*_1_ and *x*_2_ also denote the respective molecular numbers):

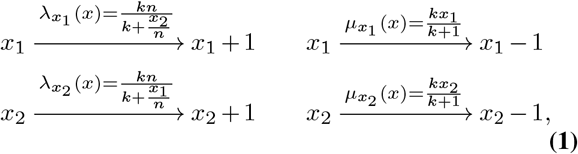

where, species *x*_2_ inhibits the production of *x*_1_ by reducing its birth rate 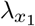 and vice versa via standard Michaelis-Menten dynamics. Both species catalyze their own degradation with death rates 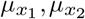. The dynamics of these reactions are governed by reaction propensities (rates) that depend on the concentration of the different biochemical species *x*_1_ and *x*_2_. In the limit of small numbers, this corresponds to a stochastic birth-death process operating on discrete (integer) numbers of molecules. We adopt this modeling framework throughout the paper and use Gillespie’s stochastic simulation algorithm (18) for all the simulations (see also SI Appendix Section A.2).

**Fig. 1.**
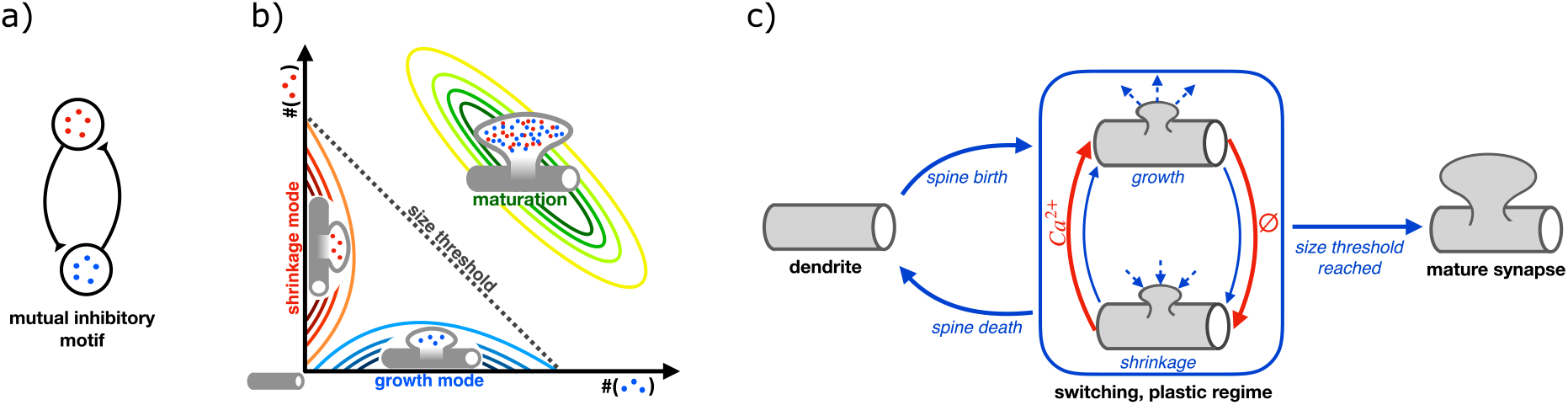
a) Illustration of mutually inhibitory connections between the two species that exhibits switch-like behavior. (b) Illustration of the dual behavior. For small system size *n*, there are two regions of attraction that result in a switching behavior between a growth (“on”) and a shrinkage (“off”) mode; for large system size, there is only one region of attraction and thus the system behaves stably. (c) Block diagram of the proposed mechanism that exploits the dual behavior for regulating plasticity.

This system is derived from the well-known toggle switch, originally used as a model of mutually repressing gene transcripts in genetically engineered E. coli (14, 17). These original studies assumed there is steep cooperativity in the birth rates of each species, resulting in two stable equilibria in the ordinary differential equations that describe the macroscopic, mean-field behavior in this system (also known as mass-action kinetics). However, the key difference between system described in Eq. (1) and the toggle switch is that Eq. (1) has a single stable equilibrium at 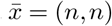 in the deterministic (large system size, *n*) limit and thus does not exhibit macroscopic switch-like behavior. We can see this immediately by calculating the so-called mean-field representation of Eq. (1), obtaining a deterministic (continuous-time and continuous-space) system of differential equations,

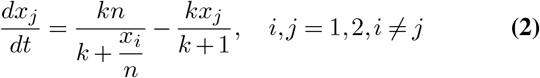

Crucially, in the small system size with a total count of *x*_*i*_ in the tens, the system dwells in two distinct modes close to the axes (see Figure 1 (b)) (15, 16). As *n* increases, however, a third mode appears close to the deterministic equilibrium that gradually absorbs the other two modes. We thus see that in this very simple system, there is a size-dependent threshold at which the system transitions between a switching and a stable behavior.

To be more precise, the stationary probability distribution (SI Appendix Section A.1) *P*^*s*^(*x*) of system Eq. (1) has one, two or three modes depending on the parameters *n* and *k*. When *n* is large, *P*^*s*^(*x*) has one mode with its peak being around 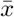. This means that 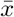 is an attractor point around which the process fluctuates. When *n* is small, *P*^*s*^(*x*) is bimodal where the two mode peaks lie along the two axes. This distribution describes a switching behavior of the system Eq. (1) between the two modes. In between these two regimes for *n, P*^*s*^(*x*) has three modes with the peaks consisting of the combination of the ones peaking along the axes and around 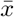 (see also SI Appendix, Figure 8). This size dependent modality happens due to transient extinctions of one of the species that becomes a prevailing phenomenon in the low number regime.

Our key insight, which we will illustrate in this simple system, is that biological systems may exploit size-dependent behavior to regulate size itself. Specifically, if the switch itself triggers growth then we have feedback between the behavior (switching or non-switching) and the system size. This constitutes a novel form of self-regulation that permits growth at small sizes, then annihilates switching behavior above a size threshold. We next demonstrate this idea with a simple model of synaptic potentiation and growth that allows the strength of an external signal to determine whether a small, immature synapse will grow and stabilize.

## Synapse size as a regulator of synaptic plasticity

Synaptic transmission depends on neurotransmitter binding to receptors in small biochemical structures called postsynaptic densities. In many types of excitatory synapses, synapse strength is partly regulated by active processes that remodel the synapse and increase its size and strength in response to external signals, including strong excitatory input that elevates calcium concentration in the vicinity of the postsynaptic density. Calcium influx in turn triggers biochemical pathways that have long been hypothesized to exhibit switch-like, or threshold behavior (19, 20).

In excitatory synapses, the postsynaptic site is often compartmentalized in a spine, a small membrane protrusion with a volume typically less than 1 *μm*^3^. The development and maintenance of synaptic contacts and of spine volume is governed by a host of biochemical signaling events (12, 21), the most extensively studied being activity dependent plasticity mechanisms - long term potentiation and depression - that are coupled to growth and retraction of the synapse respectively (20, 22–25). Widespread experimental evidence points to several consistent features of excitatory plasticity and associated structural changes in dendritic spines:

- potentiation mechanisms implement a threshold on excitatory signals and accompanying calcium influx, resulting in a switch-like decision to potentiate (20, 22–25)
- potentiation (increase in receptor count) is strongly coupled to growth of the dendritic spine, resulting in a direct correlation between synapse strength and compartment size (26, 27)
- small synapses are more susceptible to being potentiated or depressed (and eliminated); as synapses increase in size and strength their capacity to potentiate further is reduced (28–32)
- the size of dendritic spines is positively correlated to the age and life expectancy of the spine (33)
- much of the signaling machinery involved in this process involves enzymes that are present in concentrations at the nM - *μ*M range. Thus, while large spines comprise volumes that can contain hundreds or thousands of enzyme molecules, nascent spines (sometimes called filopodia) likely restrict these numbers to the tens or less (12, 13).

These observations mean it is likely that synapse growth involves a transition between microscopic and macroscopic biochemical signaling. Furthermore, they imply a biochemical switching mechanism that operates reliably in the low number/high noise regime and that becomes less sensitive as the number of signaling molecules in the postsynaptic compartment increases. Several decades of experimental work has identified candidate switching networks (9, 34, 35), most notably that involving calcium-Calmodulin/CaMKII (19, 20, 35, 36). To date, the interpretation of these studies remains controversial, with some evidence suggesting an absence of bistability (37, 38), and other work indicating that switching is context-dependent (25) or that switching deactivate rapidly following dendritic spine growth (23).

We next outline a mechanism for spine evolution that captures the key phenomena enumerated above, while potentially reconciling discrepant results concerning bistability of the underlying signaling network (Figure 1). To illustrate the model architecture, we introduce an input signal to the two-species system Eq. (1) and couple the states (molecular composition) where switching occurs to growth and shrinkage processes. This provides a generic model of a biochemical switch in a cellular compartment such as a dendritic spine. As we will show, this results in a novel form of self regulation that allows the system to operate reliably as a switch in the low number regime, and which gives way to a single stable equilibrium state as size increases.

### A size-regulated switch motif

As we showed previously, the two-species system Eq. (1) has three possible modes: one around the mean-field equilibrium in the large size limit and two along the axes in small systems (see SI Appendix, Figure 8). We now introduce feedback between the modes and system size in a toy model of growth. Without loss of generality we assume that the mode along axis *x*_1_ is the “on” mode, triggering growth, and the mode on axis *x*_2_ is the “off” mode, triggering shrinkage. When the process is in the growth or shrinkage mode, the size parameter *n* increases or decreases in proportion to the time that the process spends in the corresponding mode. The third mode around the macroscopic equilibrium is assigned to be the rest mode, hence when the process is in that mode, *n* does not change. The mode boundaries are defined simply based on the concentration of the species: we say that the process *x*(*t*) = (*x*_1_(*t*), *x*_2_(*t*))^*T*^ is in growth mode at time *t* if the concentration of *x*_1_ is larger than 75%, i.e., when *x*_1_(*t*)*/*(*x*_1_(*t*) + *x*_2_(*t*)) > 0.75. By symmetry, the process is in shrinkage mode if *x*_2_(*t*)*/*(*x*_1_(*t*) + *x*_2_(*t*)) > 0.75. Otherwise, the process is in rest mode. Note that the results can easily be extended for other choices of mode boundaries.

We call this system the size-regulated switch (SRS). As described above, the modes of the SRS on the axes represent mechanisms of shrinkage and growth, that is, the size *n* of a SRS process dynamically increases or decreases depending on which mode the process is in. When *n* decreases to zero, we eliminate the synapse. As *n* increases, the modes along the axes become smaller and shift farther along the axes (see SI Appendix, Figure 8). In addition, the system gradually forms a third mode around 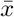 corresponding to the rest mode. Eventually, when *n* grows large enough, the system becomes unimodal with the rest mode being its only mode^1^, thus it becomes a stable system.

We next introduce two parameters to couple the shrinkage and growth modes to system size. First, we define a mode activation time threshold *t*_act_. A mode will be active and thus have an effect on the system size when the process is in that mode for longer than *t*_act_ time. Second, we use a plasticity constant *δ*_*n*_ that determines how much the system size changes when the process is in shrinkage or growth mode. When a mode is active, change in the system size is applied after every reaction in the following way: if the present time is *t* and *τ* time passed since the last reaction then the new system size is given by *n*(*t*) = *n*(*t* − *τ*)±*δ*_*n*_*τ*, where the plus and minus sign is determined by the type of mode.

For simplicity, the input to the SRS is chosen to be periodic, however, other input types are expected to lead to similar results if their role is to bias the evolution of the process towards the growth mode. To describe the periodic input, we use three parameters: (1) the period between inputs *t*_per_, (2) the duration of the input *t*_dur_ and (3) the magnitude of the input *m*_inp_. With this, the effect of the input is as follows: when *t* ∈ (*t*_per_, *t*_per_ + *t*_dur_) then the birth rate 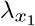 of *x*_1_ is modified to 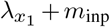.

In addition to the parameters of the mode effects and the input, the SRS is described by the initial conditions *x*(0) and *n*(0) and a constant *k* parameter that is inversely related to the inhibition strength between *x*_1_ and *x*_2_ (see Eq. (1)). Note that the shapes of the modes depend on *k* which then affects the magnitude of fluctuation in the modes, the average dwell time and the probability of switching within a time interval. For more details see SI Appendix Section C.3.

To probe the behavior of the SRS, we analyzed the survival probability and the size evolution under a range of conditions. We fixed a baseline parametrization (*x*_0_, *n*_0_, *k, t*_act_, *δ*_*n*_, *t*_per_, *t*_dur_, *m*_inp_) of the SRS and defined a set of values (values of interest) around the baseline value for each parameter 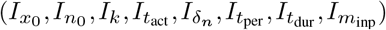 (see SI Appendix, 2 for details). To show how the parameters affect the behavior of the system, we changed the parameters one by one within their values of interest while holding the other parameters at their baseline values. For each of these parameterizations, we simulated the SRS and estimated the survival (*n*(*t*) > 0 for all *t*) probability *p*_surv_ of the system as the proportion of survived processes among all realizations, and the average system size 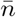 averaged over time and realizations. The results are shown in Figure 2 (c). In each plot, a point shows the proportion of the survived processes out of 10 realizations against the average system-size (averaged over time and realizations).

**Fig. 2.**
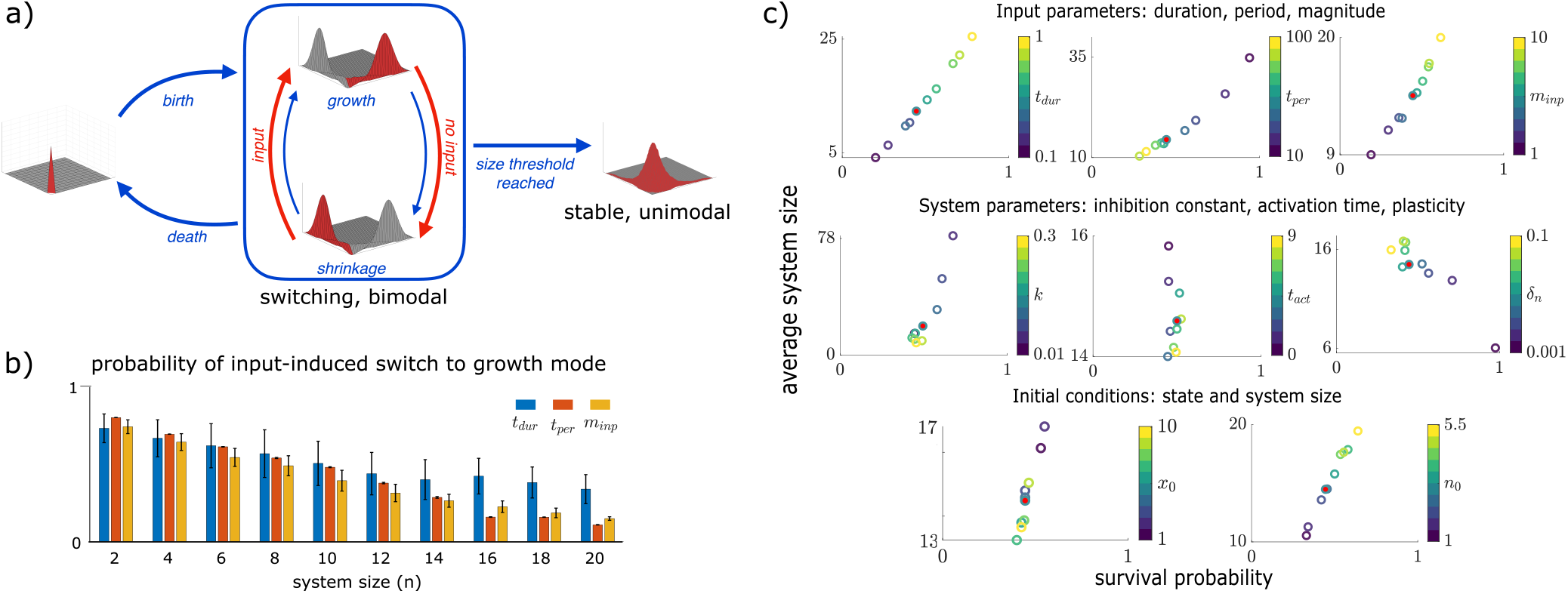
Size-regulated switch (SRS). (a) Block diagram of the SRS. (b) Demonstration of the input-sensitivity of SRS for different system sizes. As the system size increases, the SRS is less likely to switch from shrinkage mode to growth mode. This applies to inputs with a range of parameters (for details, see SI Appendix Section B). (c) Survival probability versus average system size for varying parameters. Points filled with red denote the baseline parametrization (see SI Appendix, 2). The parameters, in particular the input parameters, can tune both the average system size and the survival probability. The system parameters are less suitable for tuning survival probability than the input parameters.

The input parameters *m*_inp_, *t*_per_, *t*_dur_ affect *p*_surv_ and 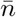 as expected: when the input is stronger, both *p*_surv_ and 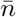 increase. Note that the input can be strengthened in three ways: it can have larger magnitude, shorter period, or longer duration. From the near linear relationship between these input features and *p*_surv_ and 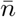, we conclude that the input can finely tune both *p*_surv_ and 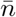.

With increasing the plasticity parameter *δ*_*n*_, the system size *n* becomes more volatile and thus reaching *n* = 0 becomes more probable. This results in a smaller *p*_surv_. However, a more volatile system induces a larger average size 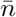 due to the bias towards growth from the applied input. In parallel, longer activation time constant *t*_act_ slightly decreases 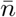 as it weakens plasticity.

The results on parameter *k* show that stronger inhibition (smaller *k*) induces larger 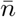 and *p*_surv_. This agrees with the observations on the system Eq. (1) that for smaller *k* parameters, the modes are farther along the axes (see SI Appendix, Figure 8) for the following reason: when modes are farther along the axes, modes are also farther apart from each other, thus switching occurs less and the input-induced bias towards growth applies more. Note that for small enough *k*, the process is likely to stay in one mode within the time scale that we simulated (see SI Appendix Figure 9).

Finally, the results on the initial conditions *x*_0_ and *n*_0_ reveal that a system with larger system size has higher survival probability and is more likely to grow bigger. The initial state *x*_0_ has a weaker but opposite effect. This is because when *x*_0_ is farther from the axes, the processes need more time to get to the modes along the axes and thus it takes longer for the system size *n* to evolve.

To sum up, the SRS provides a stochastic switching mechanism that controls system size in a manner analogous to the potentiation of dendritic spines. The sensitivity and behavior of this control mechanism can be easily tuned through parameters such as the inhibition strength (*k*) between the species and plasticity parameters (*t*_act_, *δ*_*n*_). Furthermore, external input determines system size (synapse size) and survival probability in qualitative agreement with experimental observations: smaller spines get potentiated more easily than larger spines (28–32) and the expected lifetime of smaller spines (smaller *n*_0_) is shorter (33).

## Generalization of the system

Our results so far indicate that a regulated switch can be readily implemented in a very simple and ubiquitous motif consisting of two mutually inhibitory chemical species. Such a mechanism may underlie synaptic structural plasticity and other cellular decision making processes that involve growth and signaling with small numbers of molecules. However, the specific system we have focused on is unlikely to exist in its precise form in any biological system for the simple reason that most biochemical signaling networks involve a multitude of species and exhibit diversity in their architectures and kinetic parameters. We therefore asked to what extent a size-regulated switching mechanism can generalize to multiple species and to the addition of excitatory coupling between the species.

### Two-species switching systems

To begin with, we generalize the two species system by introducing excitation between species besides inhibition and by allowing asymmetry in interaction strengths. We thus consider three types of birth-death processes *x*(*t*) = [*x*_1_(*t*), *x*_2_(*t*)]: (1) type *S*_*II*_, where the two components *x*_1_ and *x*_2_ mutually inhibit each other, (2) type *S*_*IE*_, where *x*_1_ excites *x*_2_ and *x*_2_ inhibits *x*_1_ and (3) type *S*_*EE*_, where the two components mutually excite each other. For all three cases, at any time instant, *x*_1_ and *x*_2_ increase or decrease by one in the next infinitesimal time interval with a probability proportional to the corresponding birth rates 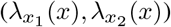 and death rates 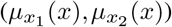. This evolution is described as follows:

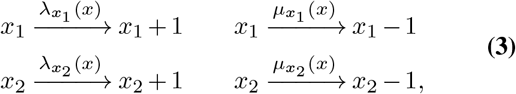

where

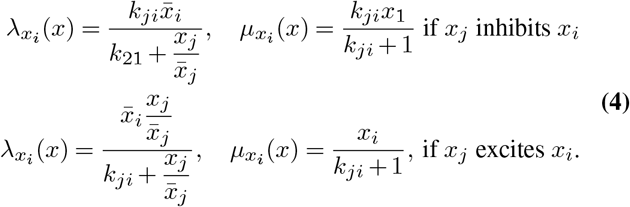

Note that in our simulations, the excitatory connections are perturbed that allows the processes to revive from the origin (see SI Appendix Section A.3).

The birth and death rates are described by the parameters *k*_12_, *k*_21_ and 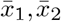. The parameters *k*_*ij*_, *i, j* = 1, 2, *i* ≠ *j* determine the strength of inhibition or excitation of *x*_*i*_ by *x*_*j*_. Analogous to the previous section, the parameters 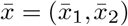 define a unique state where the birth and death rates are in balance (i.e.,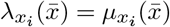 for *i* = 1, 2) providing an equilibrium of the mean-field description of Eq. (3) given by

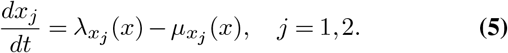

For simplicity, throughout this paper, we will assume that 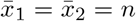 and we will call *n* the *system-size* as in previous sections. Note that the analysis can be extended to the general case. We focus on systems where 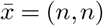 defines a unique, stable equilibrium of Eq. (5). In general, if 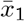 and 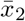 are larger than some threshold, which threshold depends on the parameters *k*_12_ and *k*_21_, the birth-death process fluctuates around 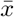 (39, Chapter 10). However, when either 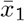 or 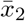 is small, other attractor points may arise. Examples of *S*_*II*_, *S*_*IE*_, and *S*_*EE*_ processes and their stationary distributions for these two regimes are illustrated in Figure 3. The columns correspond to the three types *S*_*II*_, *S*_*IE*_, and *S*_*EE*_ and the rows correspond to systems with small (top) and large (bottom) system size *n*.

**Fig. 3.**
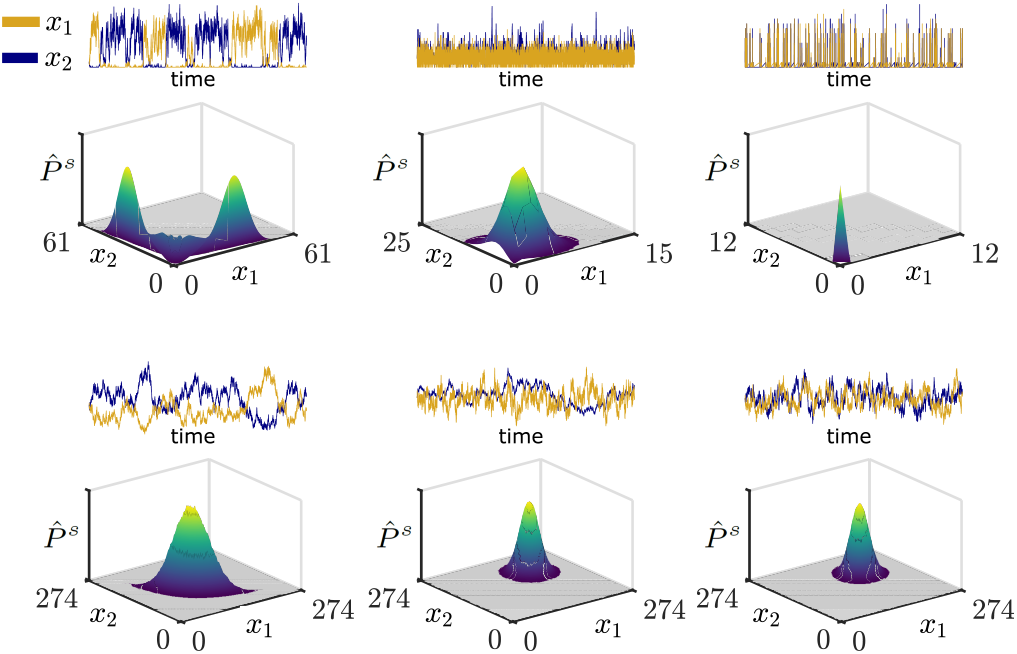
Examples of empirical stationary distributions 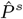 of type *S*_*II*_, *S*_*IE*_, and *S*_*EE*_ systems (from left to right, respectively) with *k*_12_ = *k*_21_ = 0.1 and small *n* = 2 (top), and large system sizes *n* = 100 (bottom). Robust multimodality appears only for *S*_*II*_ systems with small system size *n*. Segments of the corresponding processes are depicted above the distributions.

As the number of equilibrium points in deterministic systems is a key feature for describing the behavior of the system, modality of the stationary distribution is a key feature for describing the behavior of stochastic systems. A unimodal biochemical system acts like a noisy stable system and thus it is suitable for static functionality. A multimodal biochemical system on the other hand acts like a switch operator and thus it is suitable for dynamic functionality.

To understand the functionality of *S*_*II*_, *S*_*IE*_, and *S*_*EE*_ systems, we detected the parameter regimes for unimodality and multimodality by calculating the probability mass corresponding to the largest mode of the stationary probability distributions. We call this probability the Largest Mode Weight (LMW). When the LMW is close to one, the system is unimodal and when it is significantly smaller than one, the system is multimodal. For calculating the LMW, we applied our mode search algorithm (see Section A.5) that identifies the modes and the corresponding probability masses of empirical stationary probability distributions.

In Figure 4, we show the LMW of symmetric (*k*_12_ = *k*_21_, top row) and asymmetric (*k*_12_ ≠ *k*_21_, bottom row) *S*_*II*_, *S*_*IE*_, and *S*_*EE*_ systems for a range of parameters *k*_12_, *k*_21_ and *n* (see SI Appendix Section C.2 for more details on the parameters). As we can see, the *S*_*II*_ system acquired both uni-modality (yellow region) and multimodality (dark region) for a significant range of parameter triples (*k*_12_, *k*_22_, *n*). Before discussing this interesting phenomenon, we summarize the results for the other two types of systems shown in the second and third column of Figure 4: *S*_*IE*_ systems turned out to be unimodal for all parameters considered, whereas *S*_*EE*_ systems showed multimodality besides unimodality for a narrow range of parameters.

**Fig. 4.**
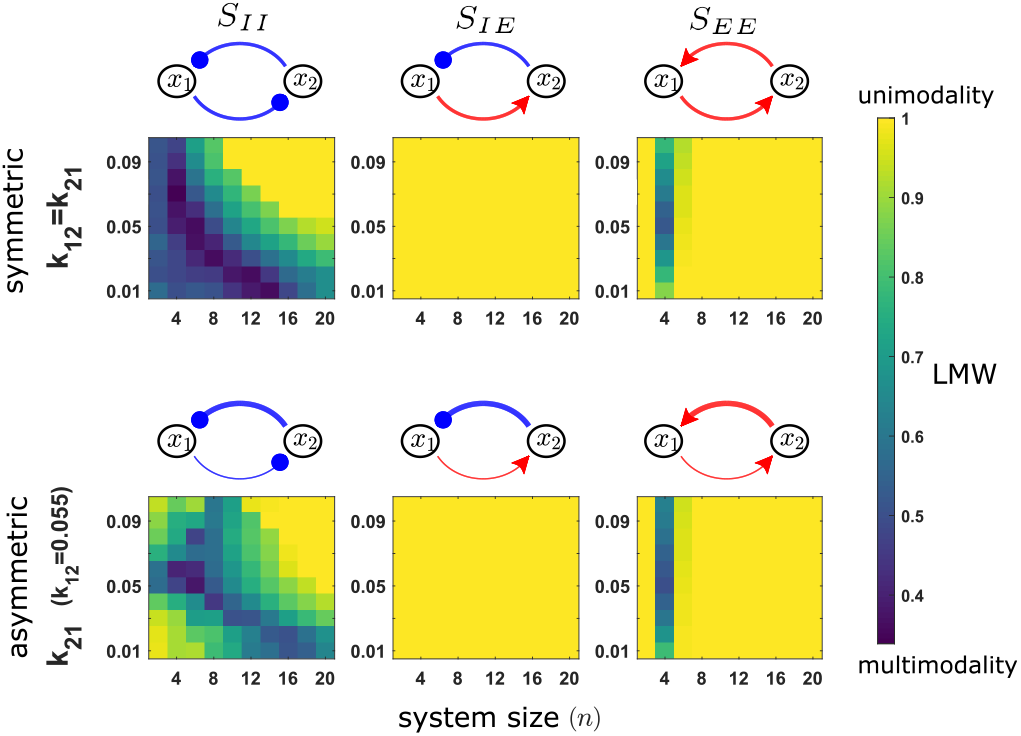
Size-dependent modality (transition from dark to yellow) typifies fully inhibitory systems (*S*_*II*_) where it is robust to asymmetry and to the strength of inhibition. This can be seen from the Largest Mode Weights (LMWs) of the empirical stationary distributions corresponding to symmetric (*k*_12_ = *k*_21_, top images) and asymmetric (*k*_12_ ≠ *k*_21_, bottom images) systems *S*_*II*_, *S*_*IE*_, and *S*_*EE*_ (from left to right). Graphs above the heatmap images illustrate the connectivities (blue: inhibition, red: excitation, thickness of the lines: connectivity strength).

This fragile multimodality of *S*_*EE*_ systems can be described as follows: The stationary distribution of *S*_*EE*_ systems has a single sharp mode at (0, 0) for small *n* representing a complete death of the species, a more spread-out single mode near 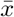 for large *n*, and a mixture of these latter modes for a few intermediate values of *n*. This type of multimodality is out of the interest of this paper for two reasons: (1) a mode corresponding to the death of the species is not suitable for dynamic functionality, e.g., it has no relevance in a feedback mechanism such as the SRS; (2) this multimodality is not robust in the parameters and thus less relevant biologically^2^.

As opposed to type *S*_*IE*_ and *S*_*EE*_ systems, when *S*_*II*_ is multimodal, its modes appear at non-trivial states, namely on the axes. In the symmetric case, when *k* = *k*_12_ = *k*_21_, the system is multimodal for small, and unimodal for large system sizes. The transition between these two happens gradually (see also SI Appendix, Figure 8) and for different system sizes depending on *k* (stronger inhibition - smaller *k* - widens the parameter regime of multimodality). In the asymmetric case, the system starts as a unimodal system (except for nearly symmetric cases), transits to multimodality for some intermediate system sizes and transits back to unimodality for large system sizes.

### Analysis of network properties that facilitate size-dependent switching

The previous results suggest that *S*_*II*_ systems have the unique feature of developing multimodality in the small system size regime that is robust for all parameters (*k*_12_, *k*_21_, *n*). To what extent does this hold more generally? We sought a means to quantify the key property of mutually inhibitory species that underlies this capacity for switching by analyzing the linearized fluctuations of the system. Such a method could open up the possibility to explore size-dependent multimodality of larger reaction networks, a possibility we will explore in three-species systems.

In general, there is no known analytic solution for stationary distributions of state-dependent nonlinear birth-death processes (15). The most accurate approximations are gained by simulating the birth-death processes and calculating the empirical distributions. However, these simulations are computationally expensive and thus do not scale well with the number of parameters (values for *k* and *n*) or with the number of species in system. Furthermore, simulations alone restrict the amount of insight that can be gleaned from results. To address this, we used an analytical approximation, the Linear Noise Approximation (LNA, (39)), that approximates the distribution up to second order. More precisely, in the LNA method, a system is approximated by a Fokker-Plank equation, whose stationary distribution is a multivariate Gaussian distribution with mean 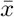 and covariance matrix Σ (see SI Appendix Section A.4). In case of multimodal distributions, this Gaussian distribution does not provide a reasonable approximation; however, as we will show, some fundamental properties of the systems, such as the direction of the fluctuations are captured by this approximation. Moreover, the Gaussian distribution approximation can give insights to the modality of the original systems through the introduction of a new measure.

We can quantify the fluctuations of birth-death processes by calculating the covariance matrix Σ from the LNA. Graphically, for a two dimensional system the covariance matrix is represented in the state-plane by an ellipse centered at the mean of the distribution. Figure 5 shows the shape of the ellipses of representative symmetric systems for *S*_*II*_, *S*_*IE*_, and *S*_*EE*_. In the *S*_*EE*_ type systems, species tend to increase or decrease simultaneously resulting in covariances with positive correlation. The ellipses representing the covariances of *S*_*IE*_ systems are close to a circle. On the other hand, *S*_*II*_ type systems have negative correlation, representing a competition between the species. When the *S*_*II*_ systems have bimodal distributions (Figure 5: values of LMW that are significantly smaller than one), the linearization exhibit fluctuations that largely exceed the positive orthant. Since the number of molecules cannot be negative, this excess tend to concentrate on the axis of the positive orthant, predicting the bimodal behavior in the original system. As the system transitions to unimodal distribution (LMW tending to one), the excess outside the positive orthant of Gaussian approximation decreases. With this observations in mind, we investigate to what extent can modality be predicted merely from the linearized system.

**Fig. 5.**
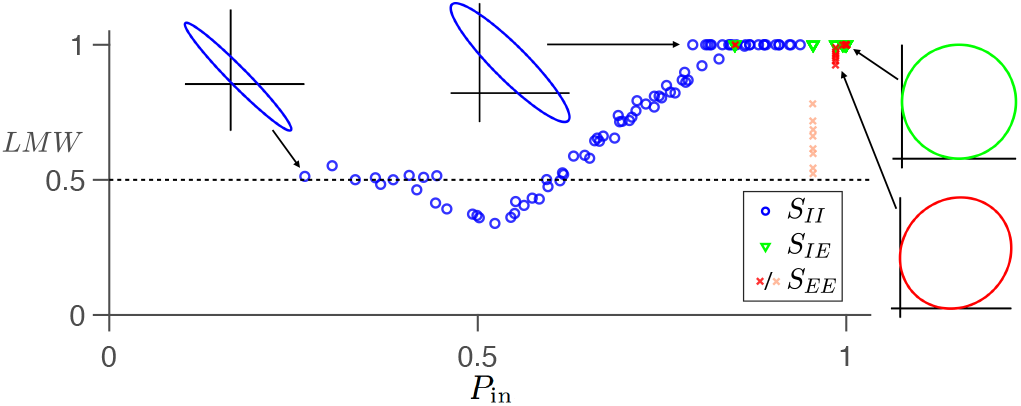
*P*_in_ provides a reliable insights of the system’s modality in the two dimensional cases. The scatter plot illustrates the relationship between the Largest Mode Weight (LMW) of the empirical distribution and the probability mass of the Gaussian distribution from the LNA inside the positive orthant (*P*_in_), for *S*_*II*_, *S*_*IE*_, and *S*_*EE*_ type systems, and parameters *k* ∈ [0.03, 0.1] and *n* ∈ [2, 20]. Note that the “weak” bimodality (see previous page) of the *S*_*EE*_ type due to the (0, 0) mode (highlighted as pink crosses) is not captured by the LNA method. The ellipses attached to the extremal points in the scatter plot represent the corresponding Gaussian approximation of the fluctuations.

To gain insight of modality prior to simulating data, we introduce a new measure using the Gaussian distribution resulted from the LNA. This measure is the probability mass of the Gaussian distribution in the positive orthant, denoted by *P*_in_. Figure 5 shows that the value of *P*_in_ is directly related to that of the LMW, and therefore to the behavior of the original system. Note that the outlier pink crosses correspond to the cases where the second mode peaks at (0, 0) and thus are out of interest of this paper.

In the next section, we investigate the modality of systems with three species. It will be shown that for higher dimensional systems, *P*_in_ can give a hint for modality and for the change of modality as *n* changes. However, it has limitations in detecting modality in the parameter *k* (see SI Appendix Section D.3). In essence, the direction of the covariance is captured well by the LNA but the magnitude of the fluctuation of multimodal systems is largely underestimated.

### Three-species switching systems

As emphasized previously, the biological relevance of our results depends on a switch being implementable in a more general reaction network. We have seen that among the two-dimensional systems that we considered, the mutual inhibitory system *S*_*II*_ robustly acquires multimodality for small system sizes. Furthermore, as the system size grows, the systems transits from multimodality to unimodality. We also showed that multi-modality can often be predicted by negative correlation in the fluctuations at the equilibrium, obtained from the linear noise approximation. We sought to test the generality of these findings in simulation. However, with the growth of the number of species, simulations quickly become intractable because both the volume of the parameter space and the support of the probability distribution of the systems grow exponentially. We therefore examined the extent to which our findings generalize from two to three dimensional systems.

Consider the time evolution of three chemical species *x*(*t*) = [*x*_1_(*t*), *x*_2_(*t*), *x*_3_(*t*)] where each species may influence the evolution of the others by inhibiting or exciting their growth. Denote the connectivity matrix between the species describing these relations by *C* ∈ {1, 0, − 1}^3×3^, where

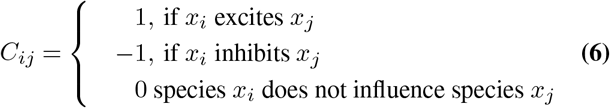

for *i, j* ∈ {1, 2, 3}, *i* ≠ *j*. Then the system evolves according to the following reaction scheme

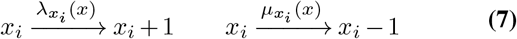

where the birth 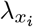 and death 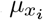 rates of a species *x*_*i*_, *i* ∈ {1, 2, 3} are given by

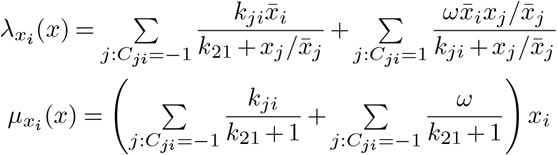

for some *ω, k*_*ij*_, and 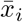 constants. Parallel to previous sections, we assume that 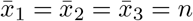 and we call *n* the system size. Note that 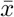 defines a unique equilibrium state of the macroscopic equations (see Eq. (5)) that guarantees the stationary distribution to be unimodal for large *n* (39, Chapter 10).^3^

Notice that for *ω* = 1, inhibition and excitation are given by the same hyperbolic functions as for the two-species systems in Eq. (4). By using the parameter *ω*, we extend our analysis with weighing the contribution of excitatory connections. We first considered seven network connectivities for three dimensional inhibitory systems (see graphs in Figure 6). These consist of all the connectivities where each species is inhibited by at least one other species and thus species do not die out indefinitely.

**Fig. 6.**
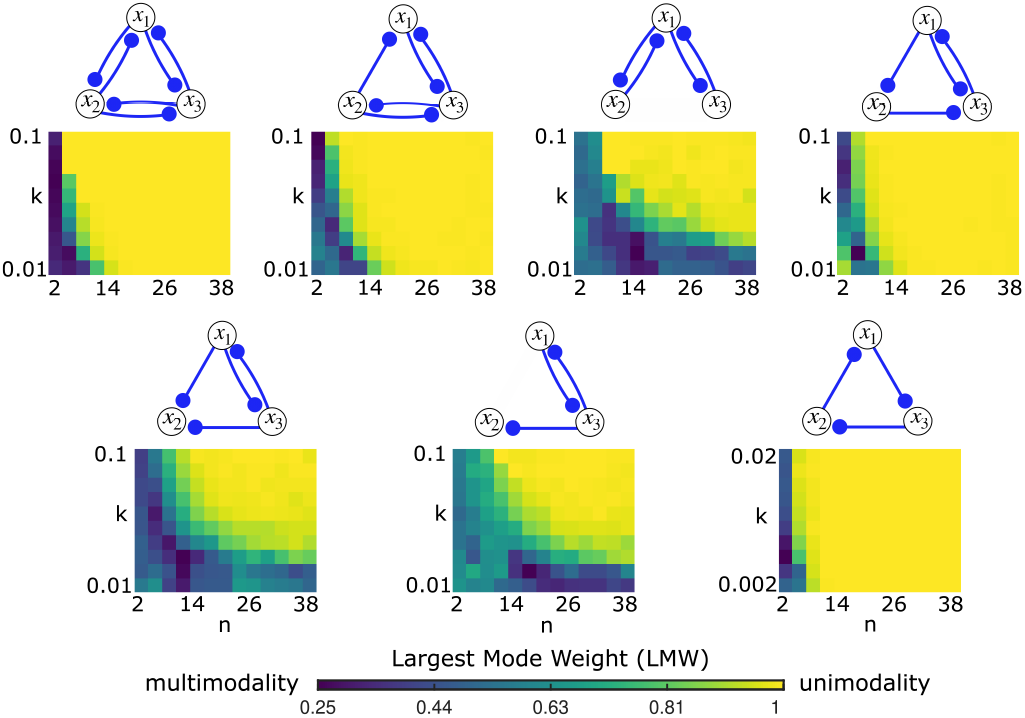
Size-dependent modality is present for all three dimensional inhibitory systems (see connectivities above the heatmap images). As the system size (*n*) increases, multimodality (dark) transitions to unimodality (yellow), where modality is measured by the LMW of the empirical distribution. This phenomenon is robust to changes in *k* (weakness of inhibition).

Both for asymmetric and symmetric inhibitory systems, (where symmetry means that *k*_*ij*_ = *k*_*lm*_ = *k* for all *i, j, l, m* ∈ {1, 2, 3}, *i* ≠ *j, l* ≠ *m*), the relationship between the system’s modality, the strength of inhibition, and the system size generalize from two dimensions to three dimensions. More precisely, multimodality is more prevailing when the system size is smaller and when the inhibition between the species is stronger. This is summarized for symmetric systems in Figure 6 by using the Largest Mode Weight (LMW) as a measure of unimodality. See also SI Appendix Figure 10 for an illustrative example. For asymmetric systems, see SI Appendix Figure 12. In Figure 6, we can see that as the system size (*n*) grows, the system transits from multimodal behavior to unimodal behavior for all the seven connectivities and this tendency is robust to the parameter *k*. Furthermore, in agreement with the two species systems, smaller *k* (stronger inhibition) widens the multimodal regime of *n*. Note that the parameter *k* shapes the stationary distribution of the system and thus *k* can be used to achieve desired separation of different modes and switching times between them.

The dependence of system modality on the parameters *k* and *n* can qualitatively be predicted by using the Linear Noise Approximation (LNA). More precisely, the trend of the Largest Mode Weight (LMW) and the probability mass inside the positive quadrant *P*_in_ calculated from the LNA agree in the parameters *k* and *n* (see Figure 11 in SI Appendix for the illustration of these findings). For quantitative prediction on the parameter regimes of multimodality, LNA is suitable if an *S*_*II*_ motif determines the modality (graphs three, five and six in Figure 6), otherwise the correspondence is qualitative. We finally extended our analysis to allow mixed excitatory and inhibitory connections. The 94 network connectivities that we obtain this way are illustrated in SI Appendix Figure 13. We used the parameter *ω* to scale the relative strengths of excitatory connections in the system. For tractability, we fixed *n* = 2 and *k*_*ij*_ = *k* = 0.01, *i, j* ∈ {1, 2, 3}, *i* ≠ *j* that provided a parametrization where all seven inhibitory systems acquired multimodality.

In the middle row of Figure 7, the LMWs of the 94 systems are presented as follows: each heatmap corresponds to one connectivity scheme (see the graphs above); within one heatmap, each row corresponds to one value of *ω* decreasing from top to bottom; within one heatmap, each column corresponds to a specific network connectivity illustrated in SI Appendix Figure 13, where these connectivities are arranged such that fully inhibitory systems gradually change into fully excitatory systems (from left to right). The number of excitatory connections of the connectivities are shown in the x-axis of the heatmaps at the bottom row.

**Fig. 7.**
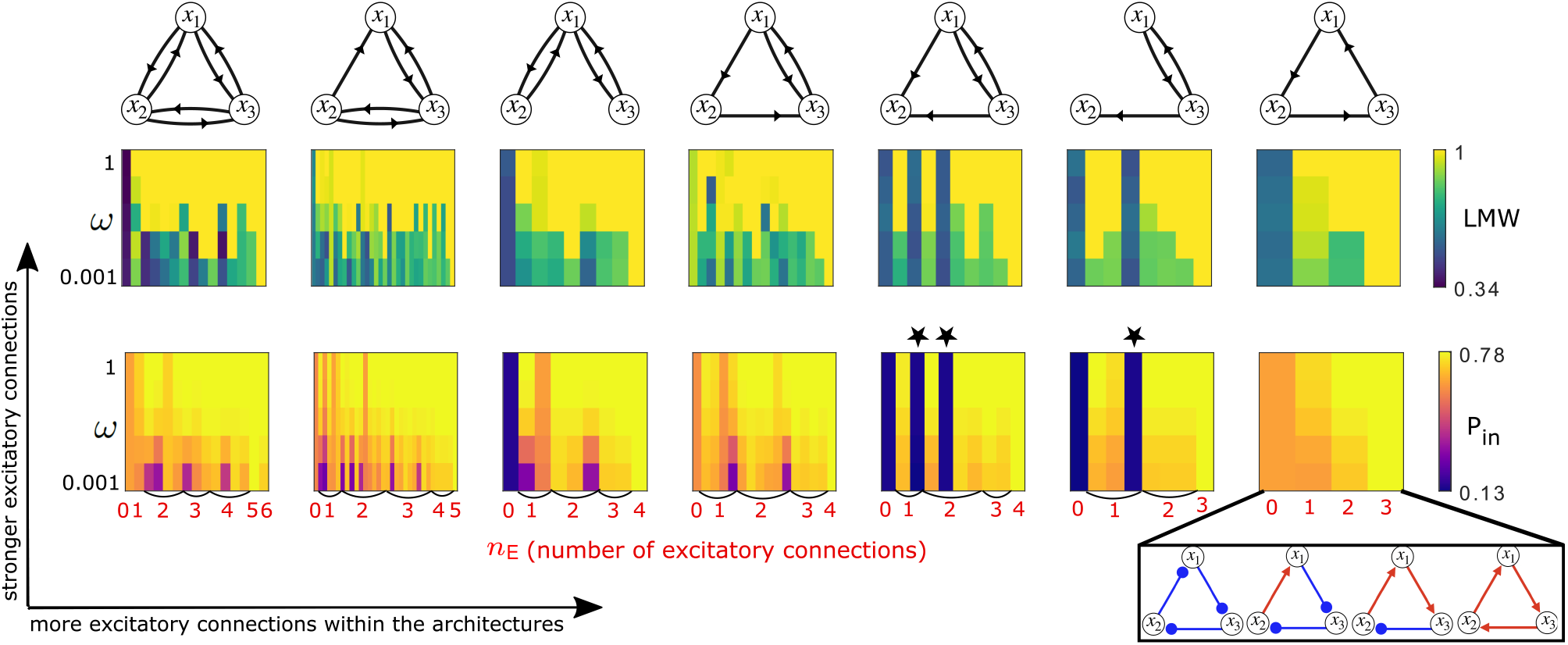
Multimodality (darker than yellow colors) is detected in systems with (1) fully inhibitory connections *n*_E_ = 0 (2) mixed connectivity where excitatory connections are weak (*ω* is small) or (3) only feedforward excitatory connections from the species with inhibitory connections (columns marked by a black star). These general tendencies can be seen from both the Largest Mode Weight (LMW) gained from empirical stationary distribution and from the probability inside the positive orthant (*P*_in_) gained from the stationary distribution of the linear noise approximation (LNA). The results are arranged by the seven connectivity schemes between the species (see graphs in top row). Within each heatmap image, rows show results for weakening contribution (*ω*) from the excitatory connections (top-to-bottom) and columns show results for different connectivities within the same scheme where from left to right, the number of excitatory connections increase (for more details, see SI Appendix, Figure 13).

**Fig. 8.**
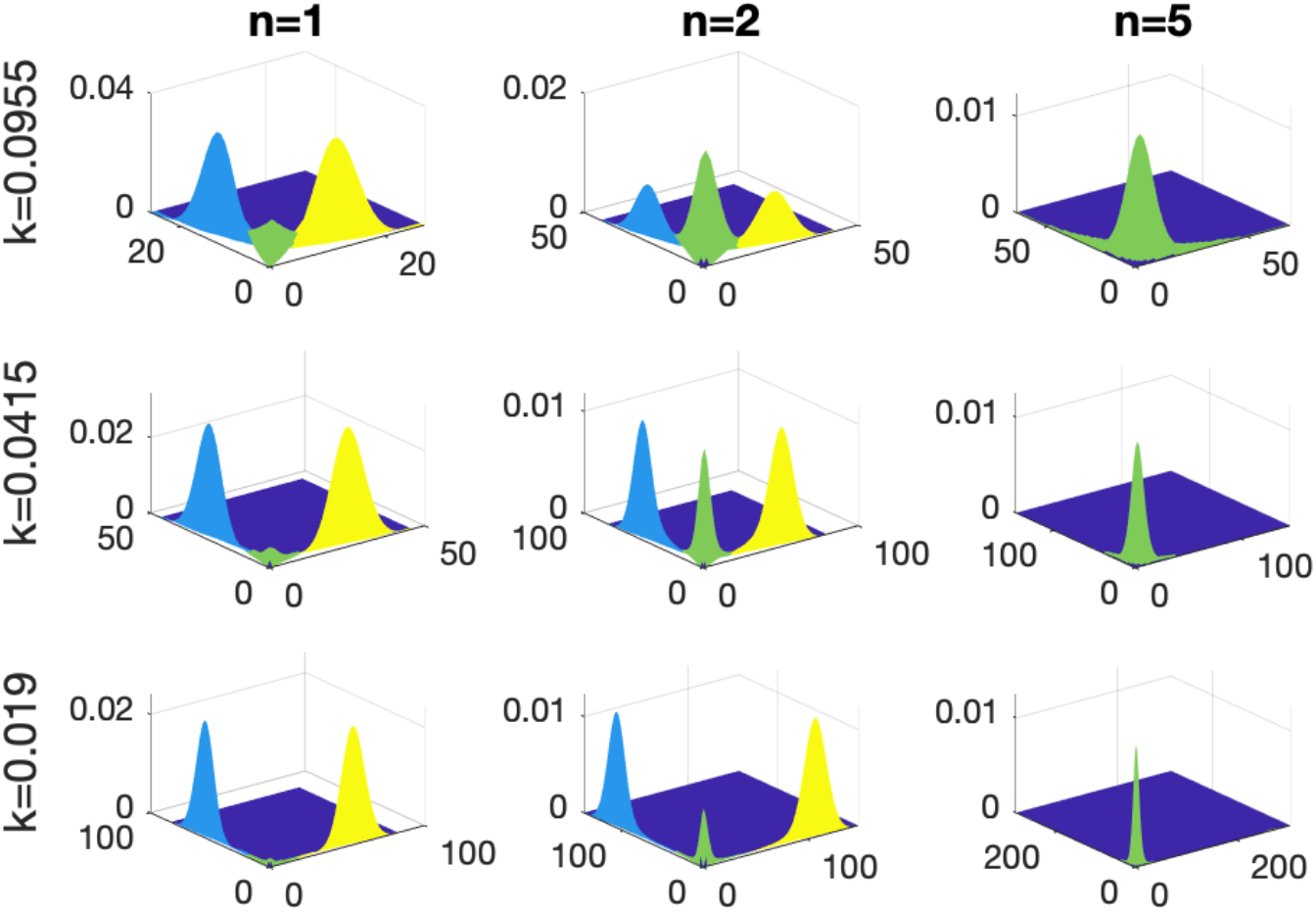
Modes of stationarity distributions of symmetric *S*_*II*_ systems as a function of system size (n) and inhibition parameter (k). The parameters are *k* = 0.0955, 0.0415, 0.019 (top-to-bottom) and *n* = 2, 6, 10 (left-to-right). The different colors represent different modes. The modes were calculated based on our mode-search algorithm described in Section A.5.

**Fig. 9.**
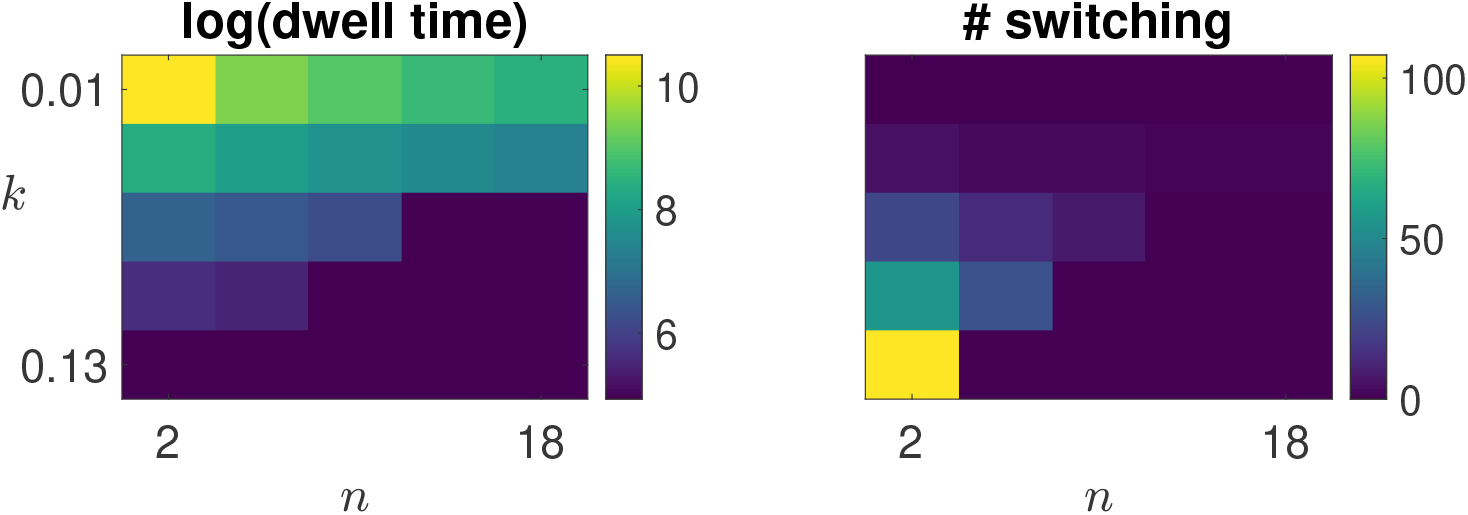
Switching properties of the symmetric *S*_*II*_ from 100 realizations of 100, 000-long (number of reactions) processes simulated using Gillespie’s SSA. Left: 10-base logarithm of dwell times for parameters *k* ∈ {0.01, 0.04, 0.07, 0.1, 0.13} and *n* ∈ {2, 6, 10, 14, 18}. Right: number of switches between the modes along the axes (for the same processes as in the left heatmap). The plots show averages over realizations, which are then (by symmetry) further averaged for the two side modes. When there was no side-mode (large *n* and *k*), both measures were set to zero.

**Fig. 10.**
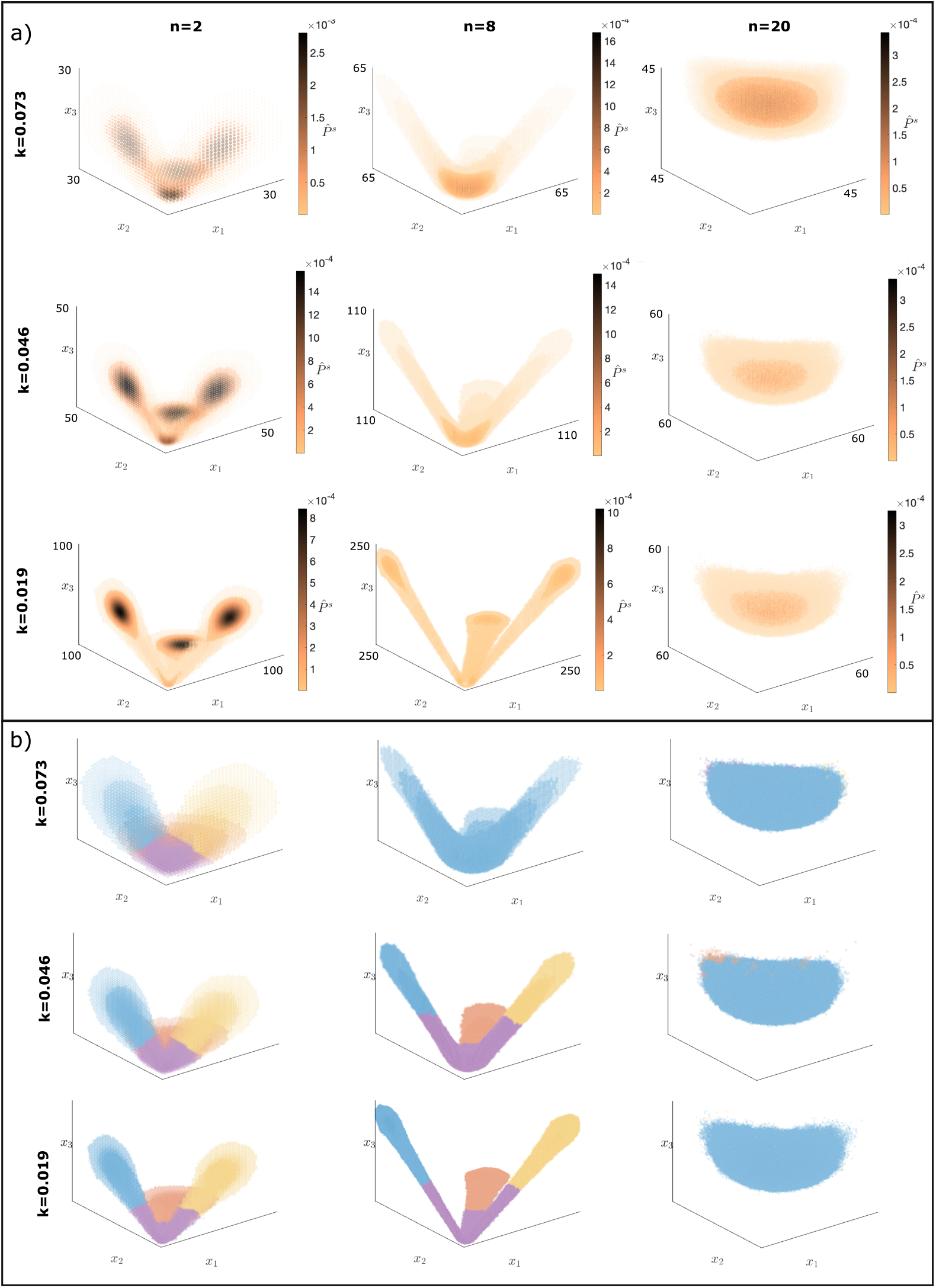
Example for three-species size dependent modality. (a) Scatter-plot of empirical stationary distributions 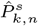 of the three-species fully connected inhibitory system (see graph on top right left of Fig. 6). The columns belong to small *n* = 2 (left), medium *n* = 8 (middle) and large *n* = 20 (right) system sizes, which illustrates the transition from multimodality to unimodality as *n* grows. The rows belong to parameters *k* = 0.073, 0.046, 0.019 (top-to-bottom). The coloring and size of a point *x* is defined by 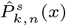. (b) Result of the mode-search algorithm (colors represent modes) of the stationary distributions 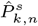 in (a).

**Fig. 11.**
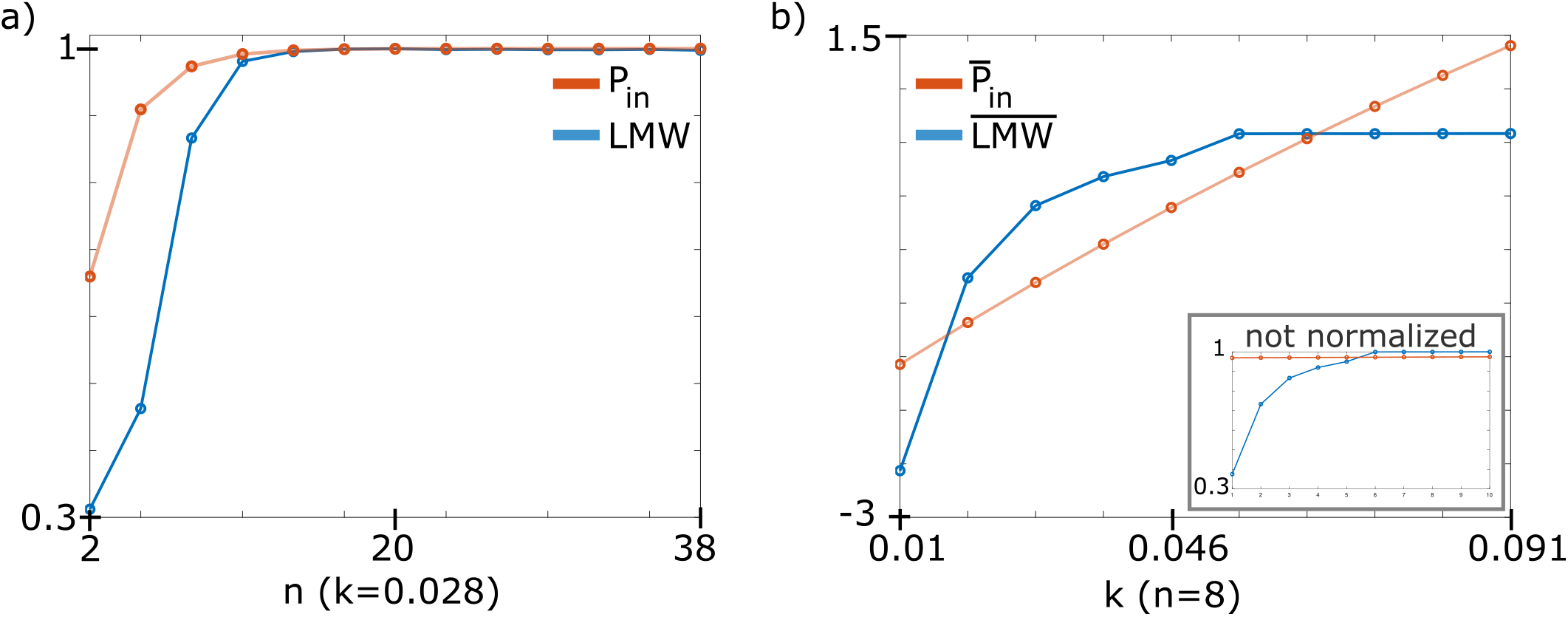
*P*_in_ and LMW of the fully inhibitory three-species system as a function of the parameters *n* and *k*. Both LMW and *P*_in_ are monotone increasing as *n* (a) and *k* (b) increases. However, *P*_in_ increases with a much smaller slope as *k* increases than the LMW. (a): *P*_in_ and LMW for *n* = 2, 5, 8, 11, 14, 17, 20, 23, 26, 29, 32, 35, 38 where *k* = 0.028 is fixed. (b): normalized 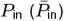 and LMW 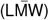 for *k* = 0.01, 0.028, 0.037, 0.046, 0.055, 0.064, 0.073, 0.082, 0.091 where *n* = 8 is fixed. In the right bottom corner, we show *P*_in_ and LMW for the same *k* and *n* values without normalization.

**Fig. 12.**
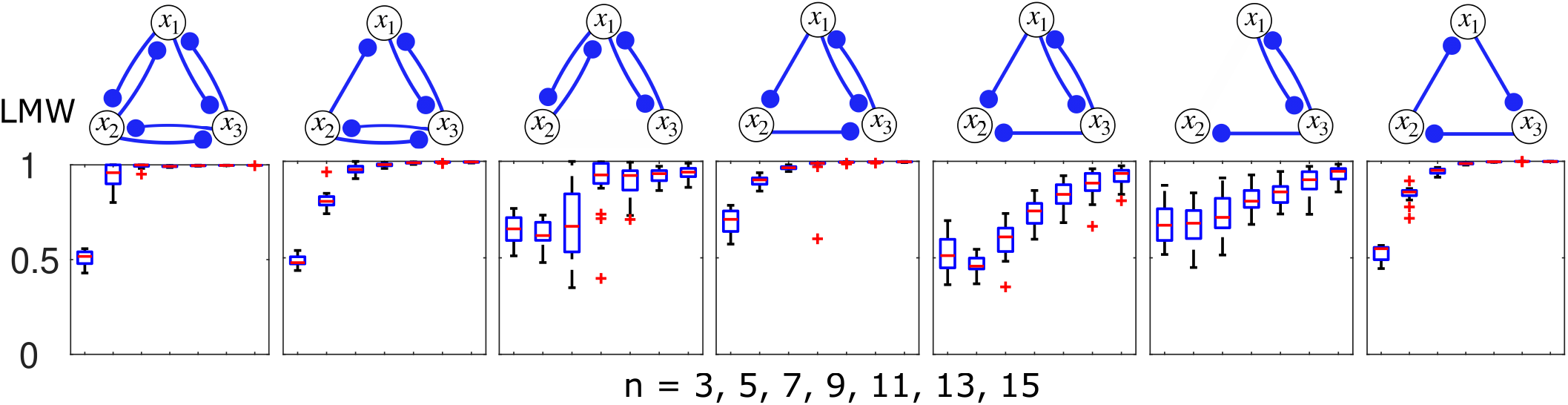
Largest mode weight (LMW) of three dimensional asymmetric inhibitory systems. For all seven connectivity schemes (top row), the statistics of LMWs from 20 randomly generated parametrization of {*k*_*ij*_}_*i,j*∈Edges_ are shown for system sizes *n* = 3, 5, 7, 9, 11, 13 and 15. The results show that, in agreement with the behavior of the symmetric inhibitory systems, the asymmetric inhibitory systems are multimodal for small *n* and as *n* increases, they gradually become unimodal.

**Fig. 13.**
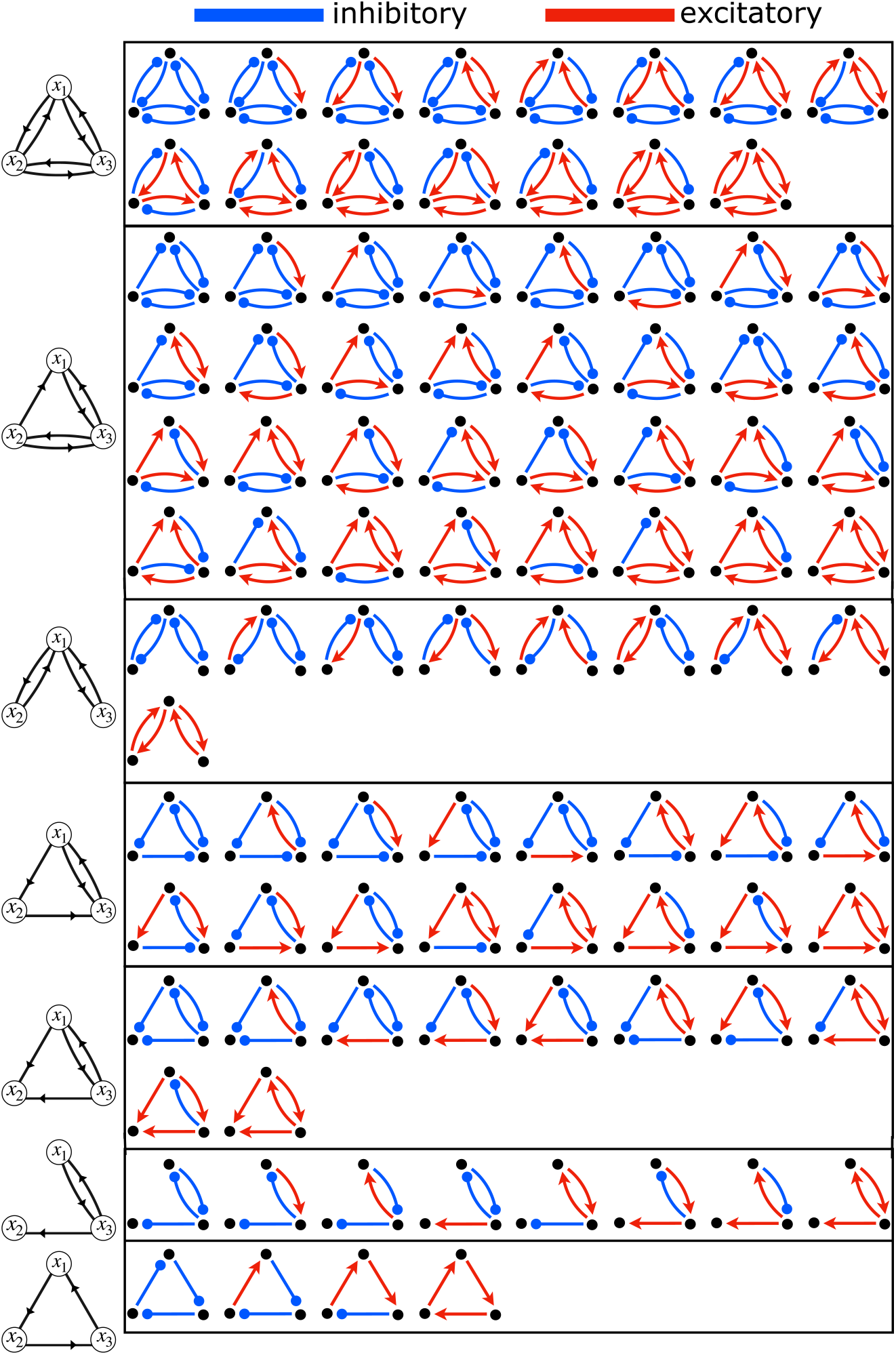
Connectivities of the three dimensional systems with inhibitory and excitatory connections. The connectivities are organized by the seven architectures that were considered in the fully inhibitory systems. The natural reading order (left-to-right and top-to-bottom) agrees with the left-to-right ordering of the connectivities in Figure 7.

The results show that if there is at least one inhibitory connection and the contribution of the excitatory connections is weak (*ω* is small), then the systems acquire multimodality. However, as the excitatory connections become stronger (*ω* is increasing), multimodal systems almost universally become unimodal. There is one category (columns marked by dark stars), when multimodality persists regardless of the strength of excitatory connections, namely, when there is an inhibitory loop between two species and there is only feedforward excitatory connection to the third species. In general, we can conclude that three dimensional systems have diverse modality if excitatory connections are feedforward or weakly feedback to an inhibitory part of the system that contains at least one loop.

In the second row of Figure 7, we present results on the probability inside the positive orthant *P*_in_ calculated from the linear noise approximation (LNA). The arrangement of the results are the same as for the LMW. These results show that *P*_in_ is a good indicator of whether a system is multimodal. However, as mentioned previously for symmetric inhibitory systems, LNA is not always suitable for determining the parameter regimes (of *ω, k* or *n*) of different modalities.

In summary, the results in Figures 6 and 7 show that system size dependent switching is not a feature of a highly specific biochemical network, and can exist in the face of significant parameter variation. The switching behavior relies on the multiple modes appearing close to the natural boundaries of biochemical networks where at least one species becomes extinct. The dependence of this phenomenon on the size is thus intuitive since extinction events are more common in small systems. Our analysis suggests that inhibitory sub-networks are necessary components for the occurrence of switching behavior because they direct fluctuations toward the natural boundaries of biochemical networks, allowing there for the formation of modes.

## Discussion

In this paper, we described and analyzed a qualitatively new kind of biochemical motif: a size-regulated stochastic switch. Our results rely on the fact that in biological compartments, where reactions occur randomly, the volume of the system can have a crucial role in shaping the behavior. Reaction networks that exhibit a stable, macroscopic equilibrium can acquire robust switching behavior in the small volume/low concentration limit. This limit typifies small synapses, where the impact of fluctuations and molecular noise has long been recognized as a potential problem for reliable signaling (40–42). Our proposal turns this problem on its head, suggesting that if breakdown of mass-action kinetics drives bimodal behavior, as happens readily in a number of simple motifs that we have analyzed, then the resulting system can function as a reliable switch. Moreover, when such a switch is coupled to growth of the compartment, a novel form of auto-regulation emerges that is consistent with the observation that many types of synapse become progressively resistant to further growth as they increase in size (28–32).

There are numerous biochemical pathways that control potentiation and growth of excitatory synapses. Perhaps the most well-known is the CamKII/PP1 pathway (35, 43–45) which mediates potentiation following strong synaptic activation and calcium influx. Classic models of this system proposed that CaMKII activates in a switch-like manner, exhibiting macroscopic bimodality in a number of detailed models of the reaction mechanism (11). Notably, the biological relevance of the parameter regime for which the CamKII concentration shows switch-like behavior has been a subject of debate (37, 46, 47). Studies (7, 11, 13) have shown that the CamKII phosphorylation can be bistable and thus fulfills the basic requirements of a switch. However, other studies (47, 48) present contradictory results. Our results provide a potential solution to this controversy by showing that the existence of bistability itself can be sensitive to the volume of space in which reactions occur, which is not a variable that is systematically accounted for in experiments.

Our results also provide more general understanding of how switches can be identified - or indeed constructed synthetically - in biochemical signaling networks. We found that inhibitory motifs play a key role in determining size-dependent switching behavior. In two and three-species systems we showed that exact stochastic simulations qualitatively agree with a general scheme where the fluctuation of mutually inhibitory systems, corresponding to negative covariance in a linear noise approximation, increase the likelihood of extinction events in small systems. Such events can lengthen the dwell time of the system in a nearby region of state space, resulting in a mode where there is no (macroscopic) equilibrium point.

In light of these results, we hypothesize that this phenomenon holds generally for inhibitory systems, regardless of dimensionality. Establishing this result without recourse to expensive simulation will require new analytical tools in nonlinear stochastic systems that thus far remain out of reach despite decades of research (15, 16, 39, 49–51). Nonetheless, the consistency of the results documented here suggests that size-dependent regulation mechanisms can readily exist in general biochemical systems. Furthermore, the robustness of switching means that global features such as survival probability and steady state volume can be tuned. We also note that the number of potential modes in the small system size regime is higher for higher dimensional systems, allowing for more complex, multi-stable switches. Given that this novel form of switching cannot be predicted by standard, continuous differential equation models that use mass action kinetics, there is the possibility that such mechanisms have been overlooked in known biochemical pathways.

More conceptually, our model is an example of how qualitative features of microscopic, discrete behavior can propagate to macroscopic scales in a controlled way. Many important biochemical reactions occur among species that are present in low copy numbers due to sub-cellular compartmentalization, small total cell volumes, or - in the case of DNA - the constraint of operating with a single copy of a molecule within a cell. As a result, cell- or tissue-wide events may be determined by microscopic, discrete, stochastic reactions. Our work provides a simple and likely common class of regulation motifs that can bridge these scales reliably.

## ACKNOWLEDGEMENTS

The authors acknowledge Andreas Petrides and Glenn Vinnicombe for their help on a preliminary form of the ideas presented here, and to Michael Rule, Thomas Burger, Mehdi Aghagolzadeh, and Dhruva Raman for technical discussions. This work was supported by ERC grant 716643 FLEXNEURO (TO) and ERC grant 670645 SWITCHLET (RS).

## A. Methods

Data and code files to reproduce the results of the paper are publicly available at https://github.com/monikajozsa/synapticswitch.

### A.1. Birth-death process and stationary distribution

A biochemical reaction network of *N* different molecular species whose interactions occur through *M* reactions can be modeled as a birth-death process (15) and represented schematically by

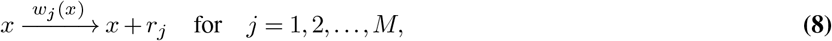

where the vector 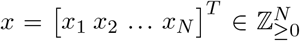 denotes the molecular count of each species in the network, and *r*_*j*_ =[*r*_1*j*_ *r*_2*j*_ … *r*_*Nj*_] ^*T*^, their variations in the numbers due to the *j*-th reaction. The coefficients of *r*_*j*_ describe a birth (*r*_*ij*_ = 1) or a death (*r*_*ij*_ = − 1) in the molecular species *i*, while *r*_*ij*_ = 0 means that the species *i* is unaffected by the *j*-th reaction. The kinetics of a reaction is modeled by the propensity function *w*_*j*_(*x*), which represents the probability per unit of time of the *j*-th reaction to occur in the network (see (50)), and it is often a function of the molecular numbers *x*. In the present work, we discriminate the propensity functions *w*_*j*_(*x*) as *λ*_*j*_(*x*) and *μ*_*j*_(*x*) for the birth and the death propensities (or rates), respectively. The Master Equation (ME) associated with the birth-death process represented in Eq. (8) is given by

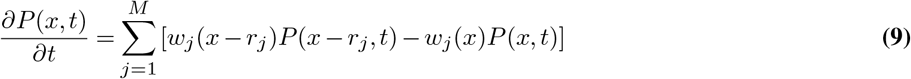

which describes the evolution of the probability distribution of the molecular numbers *x* at time *t*. Setting the time derivative in Eq. (9) to 0 gives the stationary probability distribution *P*^*s*^(*x*), which describes the probability of the molecular numbers *x* when the system reaches steady-state, i.e. for *t* → ∞ (see (39) for details).

Calculating the stationary probability distribution from Eq. (9) requires solving an infinitely dimension system of coupled equations. Therefore, in general, we cannot calculate analytic solutions of *P*^*s*^(*x*) and need to rely on estimations. In this paper, we estimate *P*^*s*^(*x*) by generating a large number of sample trajectories trough Gillespie’s algorithm (18) (see Section A.2 below).

### A.2. Gillespie algorithm

In this section, we briefly illustrate Gillespie’s Stochastic Simulation Algorithm (SSA), adopted to generate numerical realizations *x*(*t*) from the birth-death process in Eq. (8). The main idea behind the Gillespie’s algorithm is to define the conditional probability distribution *P* (*τ, j*|*x, t*), which is the probability that the next reaction will occur in the time interval [*t* + *τ, t* + *τ* + *dτ*] and this will be the *j*-th reaction, given that the number of molecules is *x* at time *t*. This probability distribution is given by

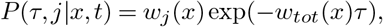

where 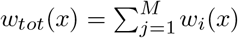. Based on this, the SSA estimates the stationary distribution as described in the following algorithm.

#### Algorithm 1 Gillespie’s SSA

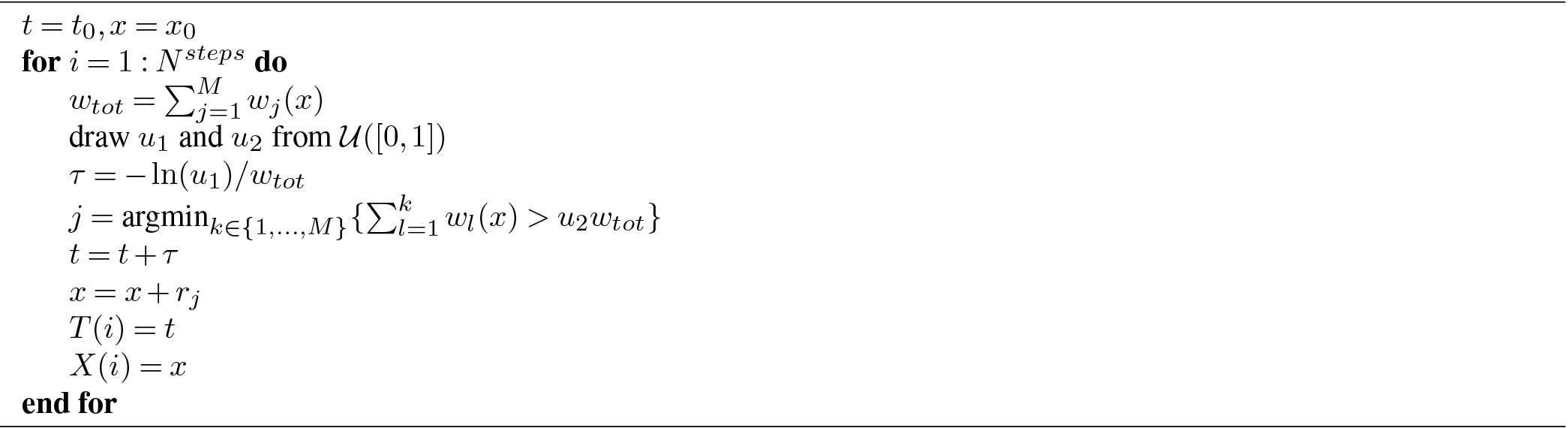

Note that in practice, we discarded the first 10% of the data in order to compensate for the bias from the initial condition *x*_0_.

### A.3. Perturbation of excitatory connections

A reaction rate that depends on excitatory connections may be zero (see Eq. (4)). This is undesirable for a stochastic system describing biological processes since such a state would act like an absorbing state and would result in an ultimate death of a species. Biological systems such as synapses and dendritic spines are not completely isolated and thus the appearance of a chemical species in a smaller compartment from a larger system such as dendritic branches is always possible. This motivates the following modeling choice: To avoid (0,0) being an absorbing state, the excitatory connections were perturbed by a fix constant *E* = 10^−4^ for all Gillespie simulations. More precisely, when *x*_*i*_ excites *x*_*j*_, the contribution of this connection to the birth of *x*_*j*_ is implemented as

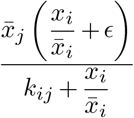

To preserve 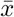 as the macroscopic equilibrium, the death rates were perturbed accordingly. For the two species systems *S*_*II*_, *S*_*IE*_ and *S*_*EE*_, the perturbed birth and death rates are summarized below in Table 1.

**Table 1.** Birth and death rates of the systems *S*_*II*_,*S*_*IE*_, and *S*_*EE*_. The parameters *k*_*ij*_, *i* ≠ *j* ∈ {1, 2} are inversely related to the strength of inhibition or excitation of *x*_*i*_ by *x*_*j*_. Due to the perturbing parameter *ϵ*, the state 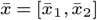 defines a unique equilibrium in the positive quadrant. The perturbing *ϵ* allows the birth of a component that is excited by the other component even when the other component is not present. In particular, in the *S*_*EE*_ system, it allows the process to revive from the state *x* = [0, 0].

| | $S_{II}$ | $S_{IE}$ | $S_{EE}$ |
| --- | --- | --- | --- |
| $\lambda_{x_1}(x)$ | $\frac{k_{21}\bar{x}_1}{k_{21} + \frac{x_2}{\bar{x}_2}}$ | $\frac{k_{21}\bar{x}_1}{k_{21} + \frac{x_2}{\bar{x}_2}}$ | $\frac{\bar{x}_1 \left( \frac{x_2}{\bar{x}_2} + \epsilon \right)}{k_{21} + \frac{x_2}{\bar{x}_2}}$ |
| $\lambda_{x_2}(x)$ | $\frac{k_{12}\bar{x}_2}{k_{12} + \frac{x_1}{\bar{x}_1}}$ | $\frac{\bar{x}_2 \left( \frac{x_1}{\bar{x}_1} + \epsilon \right)}{k_{12} + \frac{x_1}{\bar{x}_1}}$ | $\frac{\bar{x}_2 \left( \frac{x_1}{\bar{x}_1} + \epsilon \right)}{k_{12} + \frac{x_1}{\bar{x}_1}}$ |
| $\mu_{x_1}(x)$ | $\frac{k_{21}x_1}{k_{21} + 1}$ | $\frac{k_{21}x_1}{k_{21} + 1}$ | $\frac{x_1(1 + \epsilon)}{k_{21} + 1}$ |
| $\mu_{x_2}(x)$ | $\frac{k_{12}x_2}{k_{12} + 1}$ | $\frac{x_2(1 + \epsilon)}{k_{12} + 1}$ | $\frac{x_2(1 + \epsilon)}{k_{12} + 1}$ |

### A.4. Linear Noise Approximation

The Linear Noise Approximation (LNA, (39)) provides a systematic approximation of the first two moments of the solution of the Master Equation in (9). The LNA considers the Taylor expansion of the ME to the second order with respect to a parameter, called system size, which governs the size of the fluctuations. In this work, we expand the ME with respect to the parameter *n*, the macroscopic or system size, which is the equilibrium of the mean-field description associated to the birth-death process in Eq. (8). The LNA of Eq. (9) leads to a Fokker-Planck equation whose stationary solution is a multivariate Gaussian distribution with mean corresponding the mean-field equilibrium 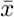 and covariance matrix Σ. This covariance matrix satisfies the Lyapunov equation

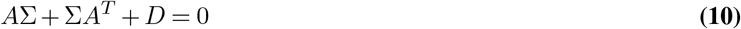

where *A* is the Jacobian matrix of the mean-field description evaluated at the equilibrium, and 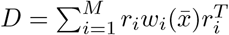 (see Section A.1 above) is the Diffusion matrix. Eq. (10) is often known in the literature as the Fluctuation-Dissipation theorem (15).

As an example, we apply the LNA method to the the three types of systems *S*_*II*_, *S*_*IE*_ and *S*_*EE*_, whose rates are described in Table 1, and we consider the symmetric case where 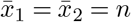 and *k*_12_ = *k*_21_, and for *ϵ* → 0. Solving Eq. (10) for each system, gives the following covariance matrices:

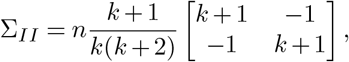

whose eigenvalues are 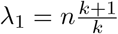 and 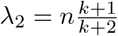, with corresponding eigenvectors *v*_1_ = [− 1, 1]^*T*^ and *v*_2_ = [1, 1]^*T*^. The covariance matrix of *S*_*II*_ highlights the negative correlation between the two species.

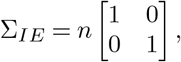

whose eigenvalues are *λ*_1_ = *n* and *λ*_2_ = *n*, with corresponding eigenvectors *v*_1_ = [1, 0]^*T*^ and *v*_2_ = [0, 1]^*T*^. The covariance matrix of *S*_*IE*_ shapes a circle with radius depending on the system-size *n*.

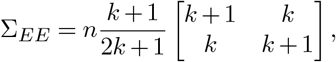

whose eigenvalues are *λ*_1_ = *n*(*k* + 1) and 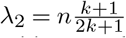, with corresponding eigenvectors *v*_1_ = [1, 1]^*T*^ and *v*_2_ = [−1, 1]^*T*^. The covariance matrix of *S*_*EE*_ highlights the positive correlation between the two species.

### A.5. Mode Search Algorithm

In order to introduce the Mode Search Algorithm applied in this paper, we first need to introduce three features of a state *x* that represents the number of molecules form each species. The first one is the empirical stationary probability density at *x*, that is 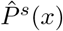. The second one is the Euclidean distance between *x* and the closest larger point *cl*(*x*) from *x* denoted by 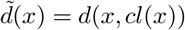 (see (52)). The third feature is more involved and requires a separate algorithm. This feature is the smallest density value *p*^ridge^(*x*) along a ridge path between *x* and the closest larger point *cl*(*x*), see (53) for similar measures used for mode detection. The ridge path ({*r*(0), …, *r*(*T*)}) is defined iteratively. Its first point (*r*(0)) is *x*, its last point (*r*(*T*)) is *cl*(*x*) and at every intermediate point along the path, the next point (*r*(*t* + 1)) is the one having the largest probability density among the neighbors of the previous point (*r*(*t*)) that are closer to *cl*(*x*) than *r*(*t*). Note that we allow the points along the path to be diagonal neighbors. For this, we define a threshold for the maximum distance of a neighbor 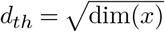 that depends on the number of species. Algorithm 2 below calculates the ridge path between between *x* and *cl*(*x*) and the smallest density value *p*^ridge^(*x*) along the path.

#### Algorithm 2 Minimum ridge path value between x and cl(x)

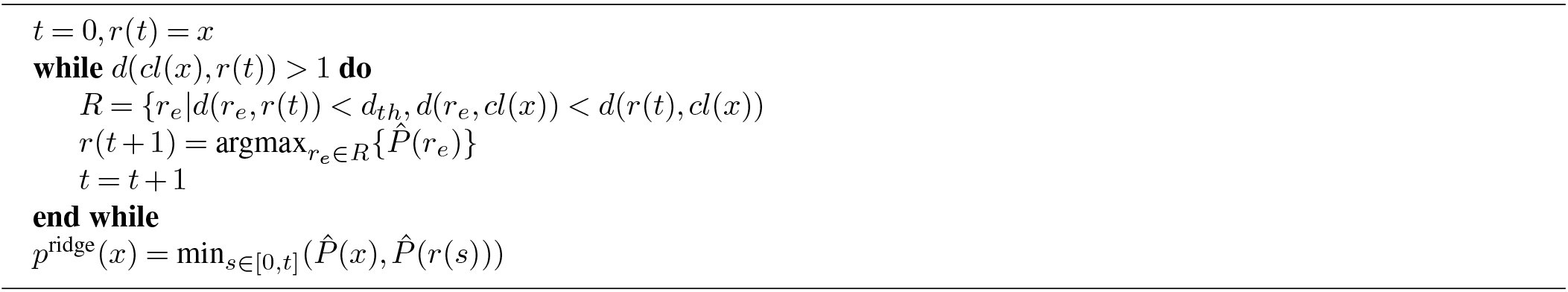

The Mode Search Algorithm assigns mode IDs to each point of the support of the empirical stationary distribution. As a first step, it detects the peaks of the modes. In principle, every local maximum is a mode peak, however, empirical stationary distributions can be rough and have spurious local maxima. In fact, the support of 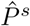 grows exponentially as the number of species and the location of the equilibrium point increase. Denote the set of local maxima by *S*. Our algorithm for selecting mode peaks from *S* is based on the three features described earlier, {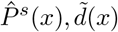, and *p*^ridge^(*x*)}_*x*∈*S*_. For each of these features and their product 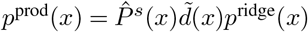, we define a threshold 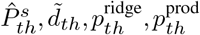. Then the subset of local maxima 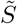 that we consider as candidate set for mode peaks is defined by

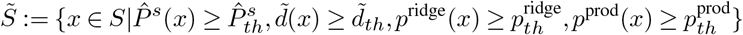

The elements of 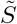 are then sorted based on their values *p*^prod^(*x*). If there are at least *N* + 1 elements, we set *N* + 1 elements with the largest *p*^*peak*^ values to be the mode peaks. Otherwise, we define all of the elements as mode peaks. Each mode peak is associated with a unique mode ID. Note that the features and thresholds are chosen such that for the global maximum *x, p*^*peak*^(*x*) = 1, 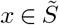. Denote the set of mode peaks by *S*^∗^.

The second step of our Mode Search Algorithm is to assign mode ID for the set of local maxima. This is described below in Algorithm 3.

For a general grid point of the support of 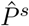, the assignment of mode ID happens based on an up-hill walk algorithm described in Algorithm 4. In summary, starting from the grid point, the algorithm iteratively steps to the neighboring point with largest 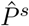 value until it arrives at a local maximum. When it arrives at a local maximum, the mode ID of the local maximum is assigned to the grid point.

#### Algorithm 3 Mode ID assignment of local maxima

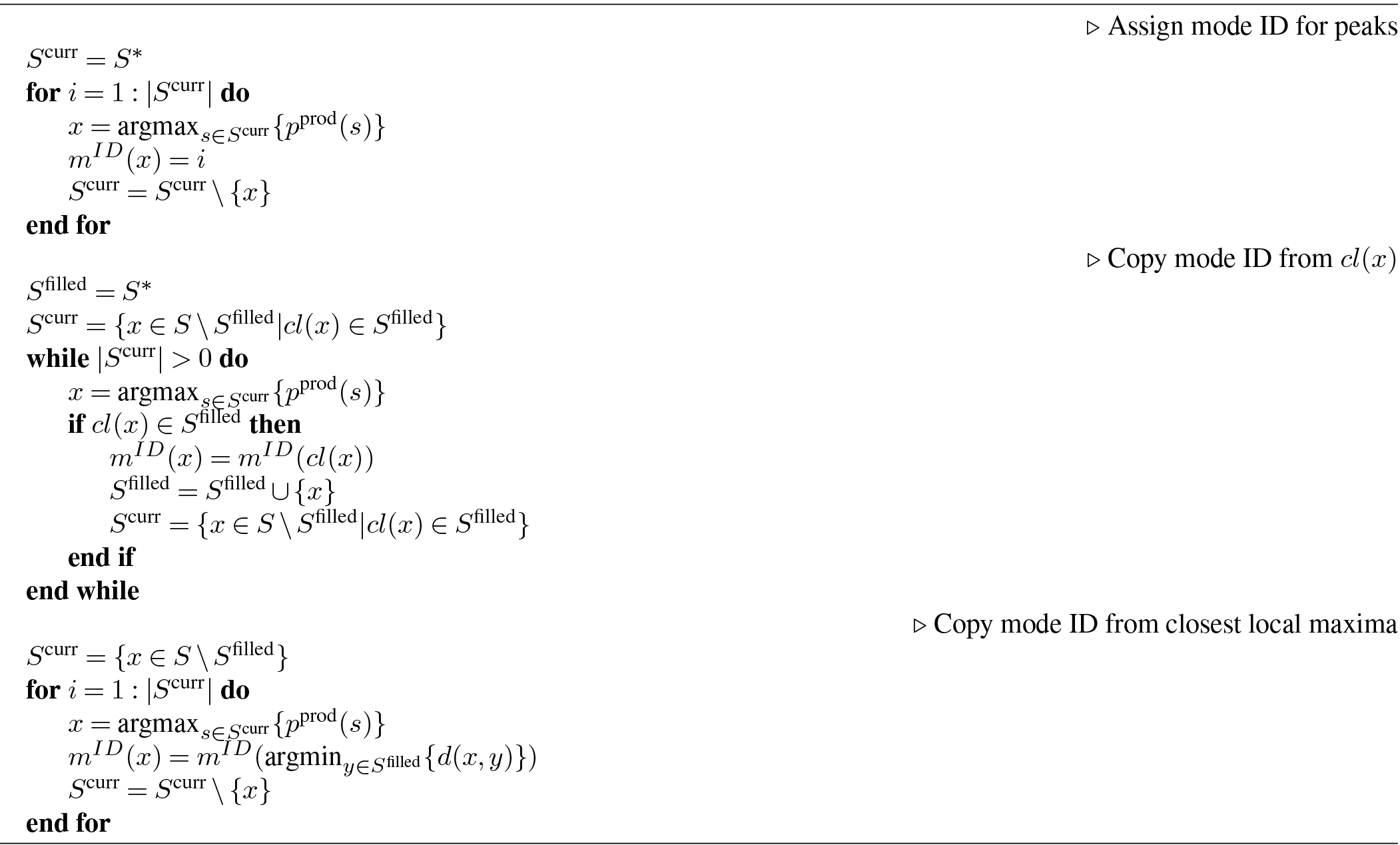

#### Algorithm 4 Mode ID assignment of all points

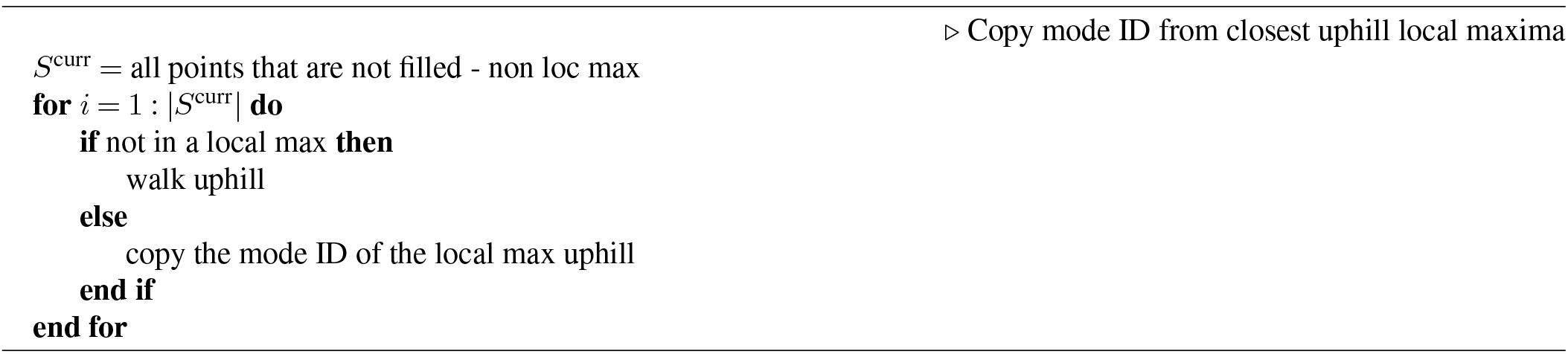

## B. Parameters of Figure 2

A key feature of the size-regulated switch (SRS) is the ability to dynamically respond to external input. This means that systems with small sizes (*n*) are more likely to be affected by the input than larger sized systems. In Figure 2 (b), we calculated the proportion of switching from shrinkage mode to growth mode following input signals with different features. We applied 28 different parameterizations of the input signals that can be categorized into three groups: (1) when *t*_per_ = 15 and *t*_dur_ = 1500 were fixed and *m*_inp_ ∈ {1, 1.5,…, 5}, (2) when *t*_dur_ = 15 and *m*_inp_ = 3 were fixed and *t*_per_ ∈ {500, 1000,…, 5000}, and (3) when *t*_per_ = 15 and *m*_inp_ = 3 were fixed and *t*_dur_ ∈ {5, 10,…, 50}.

In this analysis, we kept the system size constant throughout the simulations by setting the mode effect to be zero *δ*_*n*_ = 0 to easily handle systems with different sizes. Furthermore, we fixed *k* = 0.1389, *x*_0_ = (0, 5) and *t*_act_ = 3. For each system size (*n* = 2, 4, …, 20), we then simulated 100 realizations of 100, 000-long (number of reactions) processes.

The simulated processes were analyzed in the vicinity of the input signals. Namely, for each input pulse, a process was studied from 20 reactions prior to the first appearance of a signal to 100 reactions following the end of the input signal. We defined the condition of this segment being in shrinkage mode prior to the input signal as follows: the segment had to be in the activated shrinkage mode for the duration of at least 4 reaction intervals more than it was in the activated growth mode. Next, we defined the condition of this segment switching to growth mode following an input similarly: the segment had to be in the activated growth mode for the duration of at least 20 reaction intervals more than it was in the activated shrinkage mode. Note that the results do not depend on the exact specifications of these conditions. The reason for the asymmetry between the two conditions is that the prior condition ensures that there is a switch happening at the time of the input whereas the posterior condition ensures that the input induces a well sustained switch.

In Table 2 below, we summarize the parameters corresponding to Figure 2 (c).

**Table 2.** Parameter specifications for the simulations on size-regulated switch (SRS). For every set of parameters, we simulated 100 realizations of 100, 000 chemical reactions.

|  | parameter | symbol | baseline value | values of interest |
| --- | --- | --- | --- | --- |
| **system** | initial state | $x_0$ | $(3, 3)^T$ | fixed |
| | initial macro-size | $n_0$ | 3 | $\{1 + 0.2i\}_{i=0}^{20} \in (1, 5)$ |
| | inhibition constant | $k$ | 0.1 | $\{0.01i\}_{i=1}^{20} \in (0.01, 0.2)$ |
| **mode** | activation time threshold | $t_{\text{act}}$ | 5 | $\{0, 1, 2, \dots, 10\}$ |
| | plasticity constant | $\delta_n$ | 0.101 | $\{0.001 + 0.05i\}_{i=0}^{20} \in (0.001, 1.001)$ |
| **input** | time duration | $t_{\text{dur}}$ | 1 | fixed |
| | time period | $t_{\text{per}}$ | 50 | $\{10, 20, 30, \dots, 100\}$ |
| | magnitude | $m_{\text{inp}}$ | 5 | $\{1, 2, 3, \dots, 10\}$ |

## C. Two species systems with mixed connectivity: *S*_*II*_, *S*_*IE*_, and *S*_*EE*_ systems

### C.1. Stationary distribution of symmetric S_II_ systems

In Figure 8 below, we illustrate the parameter dependence of the modality of symmetric *S*_*II*_ systems. We observe two phenomena: (1) as the inhibition becomes stronger (*k* becomes smaller top-to-bottom), the modes become more separated; and (2) as the system size (*n*) becomes larger (left-t-right), the multiple modes merge into one mode.

### C.2. Parameters of Figure 4

In Figure 4, we show the LMW of *S*_*II*_, *S*_*IE*_, and *S*_*EE*_ systems for a range of parameters *k*_12_, *k*_21_ and system sizes *n*. For the symmetric cases (top row), the following parameters were considered 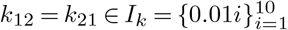. For the asymmetric cases (bottom row), *k*_12_ was fixed to a constant value of 0.055 and *k*_21_ took values from *I*_*k*_. The system size *n* took values from 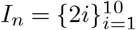. For each parametrization (*k*_12_, *k*_21_, *n*), the empirical stationary probability distribution 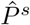 was calculated from 100 realizations of 100, 000-long (number of reactions) birth-death processes.

### C.3. Mode switching and dwell time of the symmetric S_II_ system

In this section, we describe the dependence of switching activity of the symmetric *S*_*II*_ (i.e., *k* := *k*_12_ = *k*_21_) for a range of parameters *k* and *n*. For this, we calculate the dwell times in the modes along the axes and the number of switching between them. The dwell time of a process is defined as the average time it needs to escape from one mode on the axes (starting from the peak of the mode) to the other mode on the axes (arriving at the border of the mode).

In Figure 9, we can see the dwell time and the number of switches between modes of *S*_*II*_ for different *k* and *n* parameters. The results are averaged over 100 realizations of simulations that included 100, 000 reactions. Recall that as the system size *n* increases, the growth and shrinkage modes of the size-regulated switch (SRS) at the axes become smaller and the rest mode around 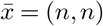 becomes larger. This is reflected in Figure 9: when the rest mode is larger, the processes visit it more easily and the dwell times in the side modes become shorter. The dwell times in the side modes eventually become zero as *n* grows and the modes on the axes gradually disappear. In other words, as *n* grows, *S*_*II*_ systems become stable and the no switching or dwell time in the modes along the axes happen. In Figure 9, we can also see that as *k* increases, the dwell time becomes shorter and the switches between modes became more frequent. This can be explained by the decreasing distance between the modes as the inhibition between the species weakens (*k* becomes larger). This tendency demonstrates that *k* can be used as a tuning parameter to achieve desired switching times.

## D. Three species inhibitory systems

### D.1. Modality of the three species inhibitory systems

The stationary distribution of three species inhibitory systems have mixed modality depending on their system size (*n*) and strength of inhibition. In Figure 10, we depict the stationary distribution of the fully connected inhibitory system for a range of parameters that illustrates the transition form multimodality to unimodality as *n* increases and the growing separation of the modes in the multimodal regime as *k* decreases.

### D.2. Parameters of Figure 6

In Figure 6, we considered system sizes 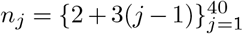 and parameters 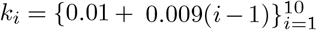 for the first six architectures and 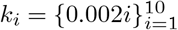 for the last one. For each system with parameter *k*_*i*_ and system size *n*_*j*_, we calculated the empirical stationary distributions 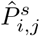 from 100 realizations of 500000-long processes. Then, we applied our mode search algorithm (see Section A.5) to identify the modes of 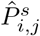. To measure uni and multimodality, we calculated the weight of the largest mode (LMW). If LMW is close to one, the system is unimodal; otherwise, it is multimodal.

### D.3. Qualitative generalization of modality in the parameters

Multimodality of three species inhibitory systems is more prevailing for smaller *k* and *n* parameters. This has been shown empirically by the Largest Mode Weight (LMW) of the stationary distribution in Figure 6. However, calculating the empirical stationary distribution and its modes is computationally expensive. In the two species system, we showed that multimodality can be detected analytically by studying the linearized fluctuations of the system, called the Linear Noise Approximation (LNA). When applied to three species systems, however, this approach proved to be only qualitatively predictive. More precisely, the probability mass inside the positive orthant *P*_in_ and the (LMW) have monotonic relationship in both *k* and *n*. However, *P*_in_ is not suitable for quantitative prediction on the parameter regimes of multimodality because the trend of *P*_in_ scales with parameter and system dependent factors. This is illustrated below in Figure 11.

### D.4. Three dimensional asymmetric inhibitory systems: the relationship between modality and system size is robust to asymmetry

Figure 12 present our results on the modality of three dimensional asymmetric inhibitory systems. We considered the same seven connectivity schemes as for the symmetric inhibitory systems and to measure modality, we calculated the Largest Mode Weight (LMW) from Gillespie SSA algorithm. In this analysis, the parameter regime of the system sizes weref chosen to be *n* = 3, 5, 7, 9, 11, 13, 15. Furthermore, the parameters *k*_*ij*_, *i, j* = 1, 2, 3, *i* ≠ *j* were generated randomly from uniform distribution *U* (0.05, 0.1) for the first six connectivity schemes and from *U* (0.005, 0.01) for the last connectivity scheme. These distributions were chosen to capture the transition of the system from multimodality to unimodality as the system size increases from 3 to 15. For each connectivity scheme and *n*, we generated 20 parameter sets {*k*_*ij*_}_*i,j*∈Edges_ independently.

### E. Three dimensional systems with inhibitory and excitatory connections

We measured the modality of three dimensional systems with mixed connectivity by the Largest Mode Weight (LMW) of the empirical stationary distributions 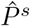. The 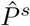 were calculated from 100 realizations of 500, 000-long processes.

In this analysis, for simplicity and tractability, we fixed the parameter of the inhibition strength *k* = 0.01 and of the system size *n* = 2. The parameter regime of the contribution of excitatory connections *ω* was chosen to be *ω* ∈ {1.0, 0.178, 0.032, 0.006, 0.001}. Below, we depict the 94 connectivities of the seven connectivity schemes that we previously introduced for fully inhibitory systems. These are the connectivities appearing in Figure 7 on the *x*-axis.

## Footnotes

1 The modes along the axes are magnitudes smaller and are far along the axes so that the process is guaranteed to stay in the rest mode for the time of interest. Note that in principle, if time is infinite, the process can visit the practically inaccessible growth and shrinkage modes and since (0, 0) is an absorbing state, the process eventually dies.

2 The sharpness of (0, 0) and the size of the intermediate *n* depend on the perturbation on the excitatory connection discussed in Table 1.

3 The excitatory connections are perturbed for the simulations similar to the two dimensional case (see SI Appendix Section A.3), which guarantees the uniqueness of equilibrium 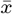.

## Bibliography

1. Thomas Eissing, Holger Conzelmann, Ernst D Gilles, Frank Allgower, Eric Bullinger, and Peter Scheurich. Bistability analyses of a caspase activation model for receptor-induced apoptosis. Journal of Biological Chemistry, 279(35):36892–36897, 2004.

2. Kenneth L Ho and Heather A Harrington. Bistability in apoptosis by receptor clustering. PLoS computational biology, 6(10):e1000956, 2010.

3. Sabrina L Spencer and Peter K Sorger. Measuring and modeling apoptosis in single cells. Cell, 144(6):926–939, 2011.

4. James E Ferrell and Eric M Machleder. The biochemical basis of an all-or-none cell fate switch in xenopus oocytes. Science, 280(5365):895–898, 1998.

5. Liang Qiao, Robert B Nachbar, Ioannis G Kevrekidis, and Stanislav Y Shvartsman. Bistability and oscillations in the huang-ferrell model of mapk signaling. PLoS computational biology, 3(9):e184, 2007.

6. Wen Xiong and James E Ferrell. A positive-feedback-based bistable ‘memory module’that governs a cell fate decision. Nature, 426(6965):460–465, 2003.

7. John E Lisman and Anatol M Zhabotinsky. A model of synaptic memory: A CaMKII/PP1 switch that potentiates transmission by organizing an AMPA receptor anchoring assembly. Neuron, 31(2):191–201, 2001. ISSN 0896-6273. doi: https://doi.org/10.1016/S0896-6273(01)00364-6.

8. Upinder S Bhalla and Ravi Iyengar. Emergent properties of networks of biological signaling pathways. Science, 283(5400):381–387, 1999.

9. Upinder S Bhalla, Prahlad T Ram, and Ravi Iyengar. Map kinase phosphatase as a locus of flexibility in a mitogen-activated protein kinase signaling network. Science, 297(5583): 1018–1023, 2002.

10. Michael Graupner and Nicolas Brunel. Stdp in a bistable synapse model based on camkii and associated signaling pathways. PLoS computational biology, 3(11):e221, 2007.

11. A M Zhabotinsky. Bistability in the Ca(2+)/calmodulin-dependent protein kinase-phosphatase system. Biophysical journal, 79(5):2211–2221, 11 2000. doi: 10.1016/S0006-3495(00)76469-1.

12. Jeanette Hellgren Kotaleski and Kim T. Blackwell. Modelling the molecular mechanisms of synaptic plasticity using systems biology approaches. Nature Reviews Neuroscience, 11 (4):239–251, 2010. doi: 10.1038/nrn2807.

13. John Lisman and Sridhar Raghavachari. Biochemical principles underlying the stable maintenance of LTP by the CaMKII/NMDAR complex. Brain Research, 1621:51–61, 2015. ISSN 0006-8993. doi: https://doi.org/10.1016/j.brainres.2014.12.010.

14. Ertugrul M Ozbudak, Mukund Thattai, Han N Lim, Boris I Shraiman, and Alexander Van Oudenaarden. Multistability in the lactose utilization network of escherichia coli. Nature, 427(6976):737–740, 2004.

15. I. Lestas, J. Paulsson, N. E. Ross, and G. Vinnicombe. Noise in gene regulatory networks. IEEE Transactions on Automatic Control, 53(Special Issue):189–200, 2008. doi: 10.1109/TAC.2007.911347.

16. Azi Lipshtat, Adiel Loinger, Nathalie Q. Balaban, and Ofer Biham. Genetic toggle switch without cooperative binding. Phys. Rev. Lett., 96:188101, May 2006.

17. Timothy S. Gardner, Charles R. Cantor, and James J. Collins. Construction of a genetic toggle switch in escherichia coli. Nature, 403(6767):339–342, 2000. doi: 10.1038/35002131.

18. Daniel T Gillespie. A general method for numerically simulating the stochastic time evolution of coupled chemical reactions. Journal of Computational Physics, 22(4):403–434, 1976. ISSN 0021-9991.

19. John E Lisman. A mechanism for memory storage insensitive to molecular turnover: a bistable autophosphorylating kinase. Proceedings of the National Academy of Sciences, 82 (9):3055–3057, 1985.

20. John Lisman, Ryohei Yasuda, and Sridhar Raghavachari. Mechanisms of CaMKII action in long-term potentiation. Nature Reviews Neuroscience, 13(3):169–182, 2012. doi: 10.1038/nrn3192.

21. Jun Nishiyama and Ryohei Yasuda. Biochemical computation for spine structural plasticity. Neuron, 87(1):63–75, 07 2015. doi: 10.1016/j.neuron.2015.05.043.

22. Jui-Yun Chang, Paula Parra-Bueno, Tal Laviv, Erzsebet M Szatmari, Seok-Jin R Lee, and Ryohei Yasuda. Camkii autophosphorylation is necessary for optimal integration of ca2+ signals during ltp induction, but not maintenance. Neuron, 94(4):800–808, 2017.

23. Seok-Jin R Lee, Yasmin Escobedo-Lozoya, Erzsebet M Szatmari, and Ryohei Yasuda. Activation of camkii in single dendritic spines during long-term potentiation. Nature, 458(7236): 299–304, 2009.

24. J Michael Bradshaw, Yoshi Kubota, Tobias Meyer, and Howard Schulman. An ultrasensitive ca2+/calmodulin-dependent protein kinase ii-protein phosphatase 1 switch facilitates specificity in postsynaptic calcium signaling. Proceedings of the National Academy of Sciences, 100(18):10512–10517, 2003.

25. Hidetoshi Urakubo, Miharu Sato, Shin Ishii, and Shinya Kuroda. In vitro reconstitution of a camkii memory switch by an nmda receptor-derived peptide. Biophysical journal, 106(6): 1414–1420, 2014.

26. Florian Engert and Tobias Bonhoeffer. Dendritic spine changes associated with hippocampal long-term synaptic plasticity. Nature, 399(6731):66–70, 1999.

27. Jon I Arellano, Ruth Benavides-Piccione, Javier DeFelipe, and Rafael Yuste. Ultrastructure of dendritic spines: correlation between synaptic and spine morphologies. Frontiers in neuroscience, 1:10, 2007.

28. Masanori Matsuzaki, Naoki Honkura, Graham C R Ellis-Davies, and Haruo Kasai. Structural basis of long-term potentiation in single dendritic spines. Nature, 429(6993):761–766, 2004. doi: 10.1038/nature02617.

29. Joshua T Trachtenberg, Brian E Chen, Graham W Knott, Guoping Feng, Joshua R Sanes, Egbert Welker, and Karel Svoboda. Long-term in vivo imaging of experience-dependent synaptic plasticity in adult cortex. Nature, 420(6917):788–794, 2002. doi: 10.1038/nature01273.

30. Cian O’Donnell, Matthew F. Nolan, and Mark C. W. van Rossum. Dendritic spine dynamics regulate the long-term stability of synaptic plasticity. Journal of Neuroscience, 31(45): 16142–16156, 2011. ISSN 0270-6474. doi: 10.1523/JNEUROSCI.2520-11.2011.

31. Anthony Holtmaat, Linda Wilbrecht, Graham W Knott, Egbert Welker, and Karel Svoboda. Experience-dependent and cell-type-specific spine growth in the neocortex. Nature, 441 (7096):979–983, 2006. doi: 10.1038/nature04783.

32. Anthony Holtmaat, Joshua Trachtenberg, Linda Wilbrecht, Gordon Shepherd, Xiaoqun Zhang, Graham Knott, and Karel Svoboda. Transient and persistent dendritic spines in the neocortex in vivo. Neuron, 45(2):279–291, 2005. doi: 10.1016/j.neuron.2005.01.003.

33. Nobuaki Yasumatsu, Masanori Matsuzaki, Takashi Miyazaki, Jun Noguchi, and Haruo Kasai. Principles of long-term dynamics of dendritic spines. Journal of Neuroscience, 28(50): 13592–13608, 2008. ISSN 0270-6474. doi: 10.1523/JNEUROSCI.0603-08.2008.

34. Jeanette Hellgren Kotaleski and Kim T Blackwell. Modelling the molecular mechanisms of synaptic plasticity using systems biology approaches. Nature Reviews Neuroscience, 11 (4):239–251, 2010.

35. John Lisman, Howard Schulman, and Hollis Cline. The molecular basis of CaMKII function in synaptic and behavioural memory. Nature Reviews Neuroscience, 3(3):175–190, 2002. doi: 10.1038/nrn753.

36. John E Lisman and Anatol M Zhabotinsky. A model of synaptic memory: a camkii/pp1 switch that potentiates transmission by organizing an ampa receptor anchoring assembly. Neuron, 31(2):191–201, 2001.

37. J.M. Bradshaw, Y Kubota, T Meyer, and H Schulman. An ultrasensitive ca2+ \calmodulindependent protein kinase ii-protein phosphatase 1 switch facilitates specificity in postsynaptic calcium signaling. Proceedings of the National Academy of Sciences of the United States of America, 100(18):10512–10517, 2003. doi: 10.1073/pnas.1932759100.

38. PJ Michalski. The delicate bistability of camkii. Biophysical journal, 105(3):794–806, 2013.

39. N. G. Van Kampen. Stochastic processes in physics and chemistry. North Holland, 2007.

40. Gabriela Antunes and Erik De Schutter. A stochastic signaling network mediates the probabilistic induction of cerebellar long-term depression. Journal of Neuroscience, 32(27): 9288–9300, 2012.

41. Daniel Choquet and Antoine Triller. The dynamic synapse. Neuron, 80(3):691–703, 2013.

42. Claire Ribrault, Ken Sekimoto, and Antoine Triller. From the stochasticity of molecular processes to the variability of synaptic transmission. Nature Reviews Neuroscience, 12(7): 375–387, 2011.

43. Yu Zhou, Eiki Takahashi, Weidong Li, Amy Halt, Brian Wiltgen, Dan Ehninger, Guo-Dong Li, Johannes W. Hell, Mary B. Kennedy, and Alcino J. Silva. Interactions between the NR2B receptor and CaMKII modulate synaptic plasticity and spatial learning. Journal of Neuroscience, 27(50):13843–13853, 2007. ISSN 0270-6474. doi: 10.1523/JNEUROSCI.4486-07.2007.

44. Seok-Jin R. Lee, Yasmin Escobedo-Lozoya, Erzsebet M. Szatmari, and Ryohei Yasuda. Activation of camkii in single dendritic spines during long-term potentiation. Nature, 458 (7236):299–304, 2009. doi: 10.1038/nature07842.

45. Salvatore Incontro, Javier Díaz-Alonso, Jillian Iafrati, Marta Vieira, Cedric S Asensio, Vikaas S Sohal, Katherine W Roche, Kevin J Bender, and Roger A Nicoll.The CaMKII/NMDA receptor complex controls hippocampal synaptic transmission by kinasedependent and independent mechanisms. Nature Communications, 9(1):2069, 2018. ISSN 2041-1723. doi: 10.1038/s41467-018-04439-7.

46. P.J. Michalski. First demonstration of bistability in camkii, a memory-related kinase. Biophysical journal, 106:1233–1235, 2014. doi: 10.1016/j.bpj.2014.01.037.

47. P.J. Michalski. The delicate bistability of CaMKII. Biophysical Journal, 105(3):794–806, 2013. ISSN 0006-3495. doi: https://doi.org/10.1016/j.bpj.2013.06.038.

48. Mariam Ordyan, Tom Bartol, Mary Kennedy, Padmini Rangamani, and Terrence Sejnowski. Interactions between calmodulin and neurogranin govern the dynamics of camkii as a leaky integrator. PLOS Computational Biology, 16(7):1–29, 07 2020. doi: 10.1371/journal.pcbi.1008015.

49. Daniel T Gillespie. The chemical langevin equation. The Journal of Chemical Physics, 113 (1):297–306, 2000.

50. Daniel T. Gillespie. Stochastic simulation of chemical kinetics. Annual Review of Physical Chemistry, 58(1):35–55, 2007. doi: 10.1146/annurev.physchem.58.032806.104637.

51. Andreas Petrides and Glenn Vinnicombe. Understanding the discrete genetic toggle switch phenomena using a discrete ‘nullcline’ construct inspired by the markov chain tree theorem. In 2017 IEEE 56th Annual Conference on Decision and Control (CDC), pages 1614–1621. IEEE, 2017.

52. Alex Rodriguez and Alessandro Laio. Clustering by fast search and find of density peaks. Science, 344(6191):1492–1496, 2014. ISSN 0036-8075. doi: 10.1126/science.1242072.

53. João C Marques and Michael B Orger. Clusterdv: a simple density-based clustering method that is robust, general and automatic. Bioinformatics, 35(12):2125–2132, 11 2018. ISSN 1367-4803. doi: 10.1093/bioinformatics/bty932.

